# Bayesian inference of tissue-migration histories in metastatic cancer from cell-lineage tracing data

**DOI:** 10.1101/2025.09.09.668633

**Authors:** Stephen J. Staklinski, Armin Scheben, Lise Brault, Rebecca Hassett, Ryan Serio, Jiawei Xing, Dawid G. Nowak, Adam Siepel

## Abstract

Cell-lineage tracing now enables direct study of tissue migration in metastatic cancer, but current reconstruction algorithms are limited by a reliance on strong parsimony assumptions and pre-estimated cell-lineage phylogenies. Here, we introduce a probabilistic modeling and inference framework, called BEAM (Bayesian Evolutionary Analysis of Metastasis), that provides richer information about complex metastatic histories. Based on the flexible BEAST 2 platform for Bayesian phylogenetics, BEAM infers a full posterior distribution over cell-lineage phylogenies and tissue migration graphs, complete with timing information. We show using simulated data that BEAM reliably outperforms current methods for inference of tissue migration graphs, especially for more complex histories. We then apply BEAM to public data sets for lung and prostate cancer, finding support for distinct modes of migration across clones and reseeding of primary tumors. Overall, BEAM serves as a powerful framework for revealing the modes, timing, and directionality of tissue migration in metastatic cancer.

## Introduction

Like all populations of proliferating cells, tumor cells adapt to their environments through an evolutionary process involving mutation, selection, and genetic drift [1]. During the past two decades, the tools of statistical molecular phylogenetics—which were originally devised to infer species trees [2–4]—have been increasingly adapted for use in reconstructing the details of tumor evolution from DNA sequences [5–8]. Recently, technologies for cell-lineage tracing and single-cell sequencing [9–13] have inspired the development of phylogenetic methods that can reconstruct lineage histories for hundreds to thousands of individual cells [14–19]. These methods have been applied not only to cancer cells but to lineage reconstruction problems in areas such as developmental biology [20–22] and neurobiology [23].

As these methods have advanced, interest in tumor lineage reconstruction has turned to the critical question of how cancer cells spread from one tissue to another through metastasis. In many cancers, metastasis represents a transition from a localized tumor that can be effectively treated with radiation or surgery to a systemic disease requiring riskier and less effective treatments such as chemotherapy and immunotherapy [24]. Several common types of cancer today—including breast, prostate, colorectal, urinary bladder, and kidney cancer—are highly treatable (with five-year survival rates >90%) when they are caught before metastasis occurs, but have dramatically poorer outcomes (with five-year survival rates of ∼40% or, in some cases, much less) after metastasis [25]. The causes of cancer mortality are complex and multifaceted, and it is an oversimplification to say that metastasis accounts for most cancer deaths [26, 27]; nevertheless metastasis is clearly one of the key events associated with the transition from treatable to untreatable disease [24].

Despite the central importance of metastasis, much remains unknown about precisely how and why it occurs. It is still not well understood whether metastases are typically initiated by single or multiple cells, whether tumor cells primarily spread through vascular or lymphatic pathways, how frequently these migrations occur, and whether they predominantly arise from the primary tumor or are dispersed widely across tissues [28–37]. In addition, it remains unclear why metastatic tumors appear preferentially in certain organs, in a manner that depends strongly on the primary cancer type and subtype, a phenomenon known as organotropism [38–42]. Strikingly, the fundamental question posed by the English surgeon Stephen Paget in his seminal article of 1889—”What is it that decides what organs shall suffer in a case of disseminated cancer?” [43]—still has no good answer 136 years hence.

As shown in several recent studies [34, 37, 44], the combination of cell-lineage tracing and phylogenetic reconstruction, typically in mouse models, promises to shed new light on these longstanding puzzles. These methods can reveal critical aspects of the rates, routes, drivers, clonality, and specific molecular changes associated with metastatic events. Notably, the primary object of interest in these studies is typically not the cell-lineage phylogeny itself but rather the induced history of migration events, which can be summarized as a “migration graph” [45]—a collapsed version of the phylogeny with nodes representing tissues and (directed) edges representing migration events between them. If both the phylogeny and the tissue-of-residence of each cell are known, then the migration graph is straightforward to derive: one simply traverses the branches of the tree and, for each branch, records a corresponding migration event if, and only if, the parent cell and child cell associated with that branch occupy different tissues. The challenge, of course, is that neither the tree nor the tissues of ancestral cells are precisely known, and uncertainty about these objects propagates to the migration graph in a complex manner. Interestingly, the problem of inferring the tissue migration graph is closely related to the problem of phylogeographic reconstruction of the movements of ancestral species; both problems require the simultaneous reconstruction of a phylogenetic tree (for cell lineages or species) and a migration graph (for tissues or geographic locations) [46–48]. In both cases, the phylogeny provides a guide but it is the induced migration history that is of primary interest.

Current methods for reconstructing tissue migration graphs generally follow a two-step process in which one or a few clonal or lineage phylogenies are first reconstructed, and then a migration history is inferred conditional on those phylogenies [45, 46, 49–51]. Typically, many possible migration histories are compatible with a given phylogeny, so a likely one is selected by maximum parsimony, i.e., by minimizing the number of tissue migrations, possibly together with other criteria. This approach represents a practical path forward and it effectively leverages powerful new tools for cell-lineage reconstruction such as Cassiopeia [14]; but it fails to exploit the tightly coupled nature of the problems of inferring the lineage tree and the migration graph. In particular, there are often many lineage trees compatible with the sequence data, particularly when mutations are sparse, but some of those trees may suggest more likely migration histories than others. Ideally, the two problems would be solved together, so that the impact on possible migration histories could be considered when inferring the lineage phylogeny (see [45] for an early attempt at such a joint solution).

A related problem is that, even when the lineage tree is fixed, many different migration histories are often plausible, and biological questions of interest are typically best addressed by considering a collection of possible histories rather than focusing on one of them [49]. For example, cancer biologists may be primarily interested in a higher-level question such as “is metastasis to bone more likely than metastasis to the liver?” or “how frequently does reseeding of the primary tumor occur?” Ideally, an analysis method would permit the investigator to evaluate the cumulative support for such a question relative to a probability distribution of possible graphs given the data, but no such method yet exists.

In this article, we address these limitations by introducing a joint probabilistic model for cell-lineage phylogenies and tissue-migration graphs, together with a procedure for Bayesian statistical inference. Our software implementation, called BEAM (Bayesian Evolutionary Analysis of Metastasis), is the first to support inference of the full posterior distribution over lineage trees, tissue migration graphs, and associated parameters given a set of DNA sequences. BEAM is implemented using the BEAST 2 platform for Bayesian phylogenetics [52], and it leverages BEAST 2’s optimized procedures for Markov chain Monte Carlo (MCMC) inference of phylogenetic parameters. In addition, BEAM supports Bayesian hypothesis testing (using MCMC-estimated Bayes factors) of any derived property of the migration graph, such as the presence of particular migration edges. We show using simulated data that BEAM accurately reconstructs features of the true migration graph over a broad range of parameters, consistently outperforming available methods. In addition, we apply BEAM to recently published lineage-tracing datasets for lung and prostate cancer models, and show that it reveals important features of the underlying migration graphs that were not evident by maximum parsimony. Overall, we show that BEAM is a promising new approach for dissecting the dynamics of metastasis across diverse cancer types.

## Results

### A joint probabilistic model for the barcode-mutation and tissue-migration processes

The problem of inferring both the cell-lineage phylogeny and tissue-migration graph can be recast as a problem of inferring a “colored” lineage tree, in which each node (cell) is assigned a color representing its tissue of residence (**Fig. 1**). Such a coloring of nodes unambiguously induces a tissue-migration graph because, as noted, migration edges correspond precisely to branches in the tree whose adjoining nodes have mismatching colors. Notice that colors (tissues) of the tips of the tree are generally known from the sampling procedure, so the problem reduces to that of inferring a likely coloring of internal nodes.

**Figure 1:**
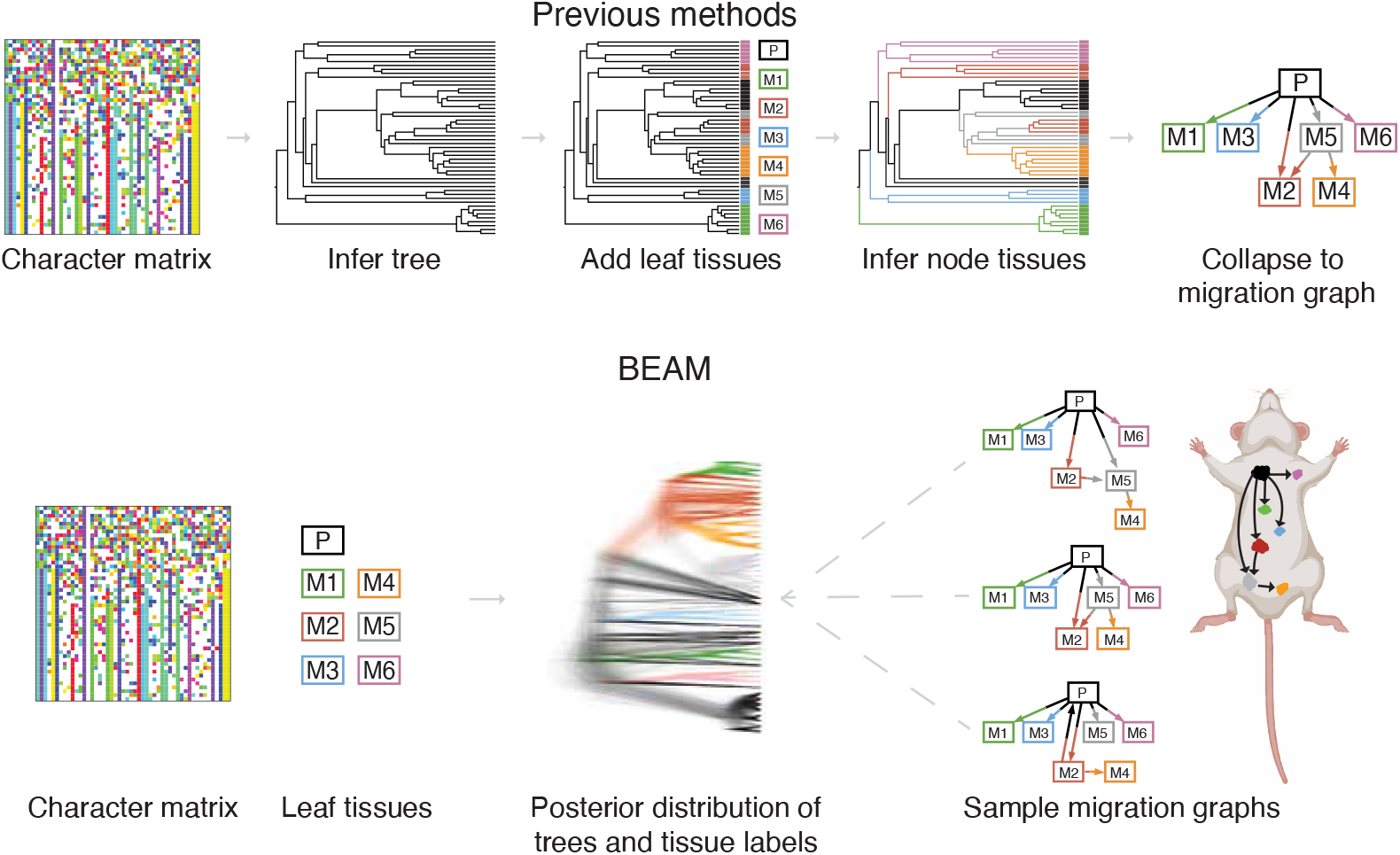
Existing methods infer a fixed lineage tree from the mutation matrix, assign observed tissue labels to the leaves of the tree, and infer ancestral tissue labels to produce one or a few migration graphs. By contrast, BEAM jointly models cell lineage trees and tissue migration histories and samples from a joint posterior distribution of trees and tissue-migration graphs. Notably, the migration graphs considered in this study are technically *directed multigraphs*, with the possibility of multiple directed edges between each pair of nodes (see **Materials & Methods**).

To define a probabilistic model for this joint process, it is therefore sufficient to extend a model for CRISPR-based lineage-tracing to allow for changes in colors (tissues), together with the accumulation of mutations, along branches of the tree. We began with a slightly simplified version of the continuous-time Markov chain (CTMC) model for CRISPR-Cas9 barcode editing implemented in TiDeTree [53] (see also [19]), and extended it with a second, conditionally independent CTMC that describes time-dependent tissue-migration events. As we will show, this second CTMC can be parameterized in a variety of ways depending on the application of interest, but our default choice is a general time reversible (GTR) parameterization.

With this model design, time-dependent transition probabilities along branches of the tree—representing joint probabilities of mutations and tissue-migrations—can be calculated as products of standard rate-matrix exponentials, and the full likelihood of the data, summing over all possible ancestral configurations, can be calculated using Felsenstein’s pruning algorithm [54] in the usual way (see **Materials & Methods**). While straightforward, this approach directly enables inference of lineage phylogenies and migration graphs together by MCMC sampling, and avoids the two-step approximation of existing methods (**Fig. 1**). We implemented this model in a software package called BEAM (Bayesian Evolutionary Analysis of Metastasis), which is built on the BEAST 2 platform for Bayesian phylogenetics [52]. An example of the output of BEAM—consisting of samples from the joint posterior distribution of lineage trees, migration graphs, timing estimates, and underlying migration rate parameters—is shown in **Suppl. Fig. S1**.

### BEAM accurately recovers tissue-migration histories from realistic simulations

To evaluate the accuracy of BEAM in reconstructing migration histories, we combined several existing simulation tools [14, 45, 55] to allow for a birth-death process at the single-cell level, migration of tumor cells between tissues, and CRISPR-editing of barcode sequences (see **Materials & Methods**). We first used this simulator to generate synthetic multi-tissue barcode sequence data in a parameter regime favorable for migration-graph reconstruction, with a relatively low migration rate (1 × 10^−6^ per cell) and a relatively high mutation rate (0.0025 mutations per barcode site per cell division). We then assessed the precision (fraction of predictions that are true) and recall (fraction of true cases that are correctly predicted) of BEAM at predicting individual edges of each simulated migration graph in 100 replicates of the simulation experiment. For comparison, we also evaluated the MACHINA [45], Metient [50], and MACH2 [51] migration-graph reconstruction methods (see **Materials & Methods** for brief descriptions). In these cases, we used pre-estimated lineage phylogenies from LAML, which currently appears to be the best method for this purpose [19]. In addition, as a baseline comparison, we assigned tissues (colors) to ancestral nodes in the same LAML-derived lineage phylogenies by simple Fitch-Hartigan parsimony [56, 57]. Finally, we trivially assigned to each ancestral node either the most common tissue label among its descendant leaves (Consensus) or a randomly selected tissue (Random).

We found that BEAM performed well in these tests, achieving nearly perfect edge-wise precision (>95%) up to a recall rate of ∼70%, after which the precision declined rapidly (**Fig. 2A**). In this parameter regime, MACHINA, Metient, and MACH2 exhibited comparable but slightly less favorable tradeoffs between precision and recall. Notably, however, MACHINA—because it produces a single reconstructed migration graph—only appears as a point on a precision/recall graph. Both Metient and MACH2 have the potential to report multiple solutions, but we found that they tended to be similar to one another, and in the case of Metient we followed its authors in reporting a single solution for performance evaluation (see **Materials & Methods**); we reported multiple solutions for MACH2 but they tended to fall in a narrow range on the graph. By contrast, BEAM permits a broad range of tradeoffs between precision and recall, by allowing the user to choose the threshold for the estimated posterior probability for each edge. Interestingly, the simple Fitch-Hartigan parsimony approach also often performed well in this regime. All methods easily outperformed the Consensus and Random baselines. For comparison with previous publications [45,50,51], we also computed the F1 score for all methods, using a posterior probability threshold of 0.5 for each edge in BEAM (**Fig. 2B**). By this metric, the rank order of methods was similar and BEAM maintained a statistically significant advantage over MACHINA, Metient, and MACH2. Examples of ground-truth migration graphs, posterior probability-weighted edge graphs, 0.5-posterior probability consensus graphs, and individual posterior distribution samples are provided in **Suppl. Fig. S2**.

**Figure 2:**
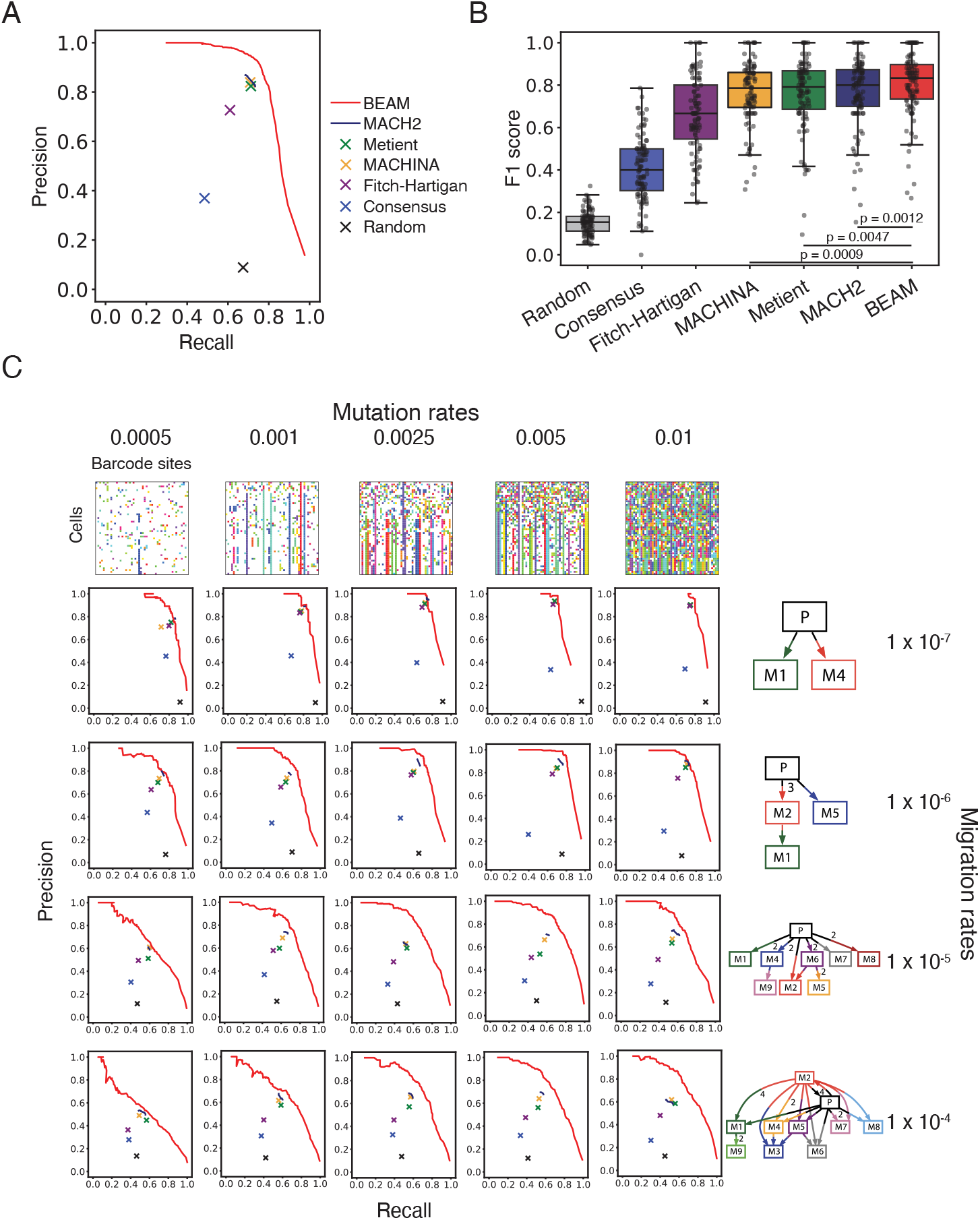
**A.** Precision-recall curves for individual edges in simulated migration graphs based on a migration rate of 1 × 10^−6^ per cell and a mutation rate of 0.0025 mutations per barcode site per cell division. The full curve for BEAM results from varying a threshold for the estimated posterior probability of each edge. Individual points are shown for solutions obtained by MACHINA [45], Metient [50], Fitch-Hartigan parsimony, and two baseline methods (Consensus and Random; see text). A curve is shown for MACH2 [51] but it is restricted in length. In all cases, mean values over 100 simulations are plotted. **B**. F1 scores for the same experiment as in panel A, based on a threshold of 0.5 posterior probability for BEAM and MACH2. Reported *p*-values are from paired *t*-tests. **C**. Precision-recall curves for the same methods under variable mutation and migration rates. Representative mutation matrices and migration graphs are shown.

To see whether these trends continued with other choices of parameters, we carried out a series of similar experiments with barcode mutation rates ranging from 0.0005 to 0.01 mutations per site per cell division and migration rates ranging from 1 × 10^−7^ to 1 × 10^−4^ per cell, executing 20 simulation replicates for each parameter combination (see **Materials & Methods**). These scenarios were designed to cover a range from cases of many mutations and few migrations, where lineage trees should be well-resolved and migration histories should be simple, to cases of few mutations and many migrations, where migration histories should be more complex and uncertainty about them should be much greater.

We found, as expected, that the prediction performance of most methods improved with both increasing mutation rate and decreasing migration rate (**Fig. 2C**). Not surprisingly, all but the baseline methods performed well at high mutation and low migration rates (top right of **Fig. 2C**), but performed considerably worse at low mutation and high migration rates (bottom left of **Fig. 2C**). Nevertheless, BEAM reliably out-performed all other methods across all parameter values. Interestingly, the performance improvement from BEAM was most pronounced at intermediate-to-high mutation and migration rates (particularly toward the bottom-right of **Fig. 2C**), indicating that the advantages of the Bayesian approach were most evident when there was ample phylogenetic signal in the data (from higher mutation rates) but substantial uncertainty about the structure of the migration graph. In this regime, the MACHINA, Metient, and MACH2 methods also showed a greater performance advantage over simple Fitch-Hartigan parsimony. When we stratified our simulated data based on features of the underlying ground-truth migration graphs, BEAM maintained performance across variable migration counts (**Suppl. Fig. S3A**), co-migration counts (**Suppl. Fig. S3B**), monoclonal or polyclonal seeding events (**Suppl. Fig. S3C**), metastatic-to-metastatic seeding events (**Suppl. Fig. S3D**), and primary reseeding events (**Suppl. Fig. S3E**). Overall, BEAM performed well across a range of migration dynamics and proved to be robust to variable levels of information content in the data.

To better understand how BEAM’s predicted migration graphs differed from those reconstructed by parsimony, we carried out a direct comparison on the same trees. We used BEAM to sample tip-labeled lineage trees from the posterior distribution, and then, for those trees, we calculated both the number of migration events predicted by BEAM and the minimum number possible according to Fitch-Hartigan parsimony [34]. The difference between these quantities is a measure of the excess migrations predicted by BEAM. We found, not surprisingly, that in favorable regimes for migration graph reconstruction, this excess was low, with values of zero in most cases and occasional deviations of 1–3 migrations (**Suppl. Fig. S4A**). With lower mutation rates or higher migration rates, however, we observed larger excesses, with the BEAM predictions often having as many as 5, and sometimes 15 or more events than the minimum possible (**Suppl. Fig. S4B-C**). Given the clear performance advantage of BEAM in this regime (**Fig. 2C**), this observation implies that parsimony often oversimplifies the true history and BEAM gets closer to the truth by allowing for violations of strong parsimony assumptions, particularly along longer branches of the tree. Altogether, our analysis of simulated data shows that BEAM reliably infers diverse migration histories, appropriately manages uncertainty, and allows the data to guide solutions toward or away from parsimony solutions.

### BEAM reveals complex patterns of tissue migration in lung and prostate cancer models

Having demonstrated that BEAM performed well on simulated data, we turned to real data from two recent applications of lineage-tracing in mouse models of metastasis, one in lung cancer [34] and one in prostate cancer [37]. Despite numerous differences in their strategies for cancer initiation (an orthotopic xenograft model of lung cancer in immunodeficient mice vs. a somatically engineered mouse model [SEMM] of prostate cancer in immunocompetent mice) and barcode sequencing (single-cell RNA sequencing vs. bulk sequencing of PCR-amplified DNA), these studies both produced mutation matrices representing many barcoded cells from distinct tissues. In both cases, cells were grouped by barcode similarity into clonal populations (CPs) representing lineage trees from distinct founder cells.

The two datasets had previously been analyzed using different parsimony strategies (Fitch-Hartigan parsimony in [34] and MACHINA in [37]). For a uniform comparison point for BEAM, we re-analyzed them using current best practices, including LAML [19] (instead of Cassiopeia [14]) for lineage-tree reconstruction and MACH2 [51] for migration graph inference. The tissue migration graphs reconstructed by BEAM and MACH2 were broadly similar with somewhat more complexity evident in the BEAM reconstructions on average, consistent with our simulation results. For example, in the 0.5-posterior-probability consensus graphs across lung cancer CPs, MACH2 averaged 12.8 migrations and 3.5 co-migrations, while BEAM av-eraged 13.9 migrations and 4.6 co-migrations; across prostate cancer CPs, MACH2 averaged 8.2 migrations and 2.8 co-migrations, compared to BEAM averages of 16.6 migrations and 5.4 co-migrations. When we examined individual CPs, we found that, in some cases, the two methods predicted quite different evolutionary histories (e.g., **Suppl. Fig. S5**), but in other cases the predictions were similar (e.g., **Suppl. Fig. S6**). The CPs tended to vary considerably in size, mutation content, and migration graph complexity between the lung cancer and prostate cancer datasets, providing a range of scenarios in which to evaluate BEAM.

A key open question is whether migration events predominantly arise from the primary tumor, or whether, by contrast, migrations occur at appreciable frequencies between metastatic sites or from metastatic sites back to the primary tumor. The original analysis of the lung cancer data [34] reported relatively high rates of both metastasis-to-metastasis (M2M) and primary reseeding (PR) events, with ∼90% of CPs displaying M2M events and ∼65% of CPs displaying PR events. BEAM and MACH2 were reasonably consistent with the previous analysis overall, but with somewhat lower estimates of the incidence of PR events: MACH2 reported ∼95% M2M and ∼50% PR, and, at a posterior probability threshold of 0.8, BEAM reported ∼94% and ∼45% respectively (**Fig. 3A**). Notably, however, the BEAM predictions were quite sensitive to the edgewise posterior-probability threshold, with the rate of M2M events varying from ∼96% at a threshold of 0.5 to ∼66% at a threshold of 0.99, and the rate of PR events varying from ∼53% to ∼13%. MACH2 typically produced a small set of equally parsimonious graphs that tended to be highly similar to one another, making the results largely insensitive to the choice of edgewise posterior-probability threshold. Overall, our Bayesian framework appears to assign high uncertainty to these secondary migration events, which generally occur later in tumor development and have weaker support in the mutation matrix. In particular, our most conservative estimates of the rate of PR events, at ∼13–23%, are several times lower than the previous estimate of ∼65%. Nevertheless, our analysis supports that both M2M and PR events occur with non-negligible frequency.

**Figure 3:**
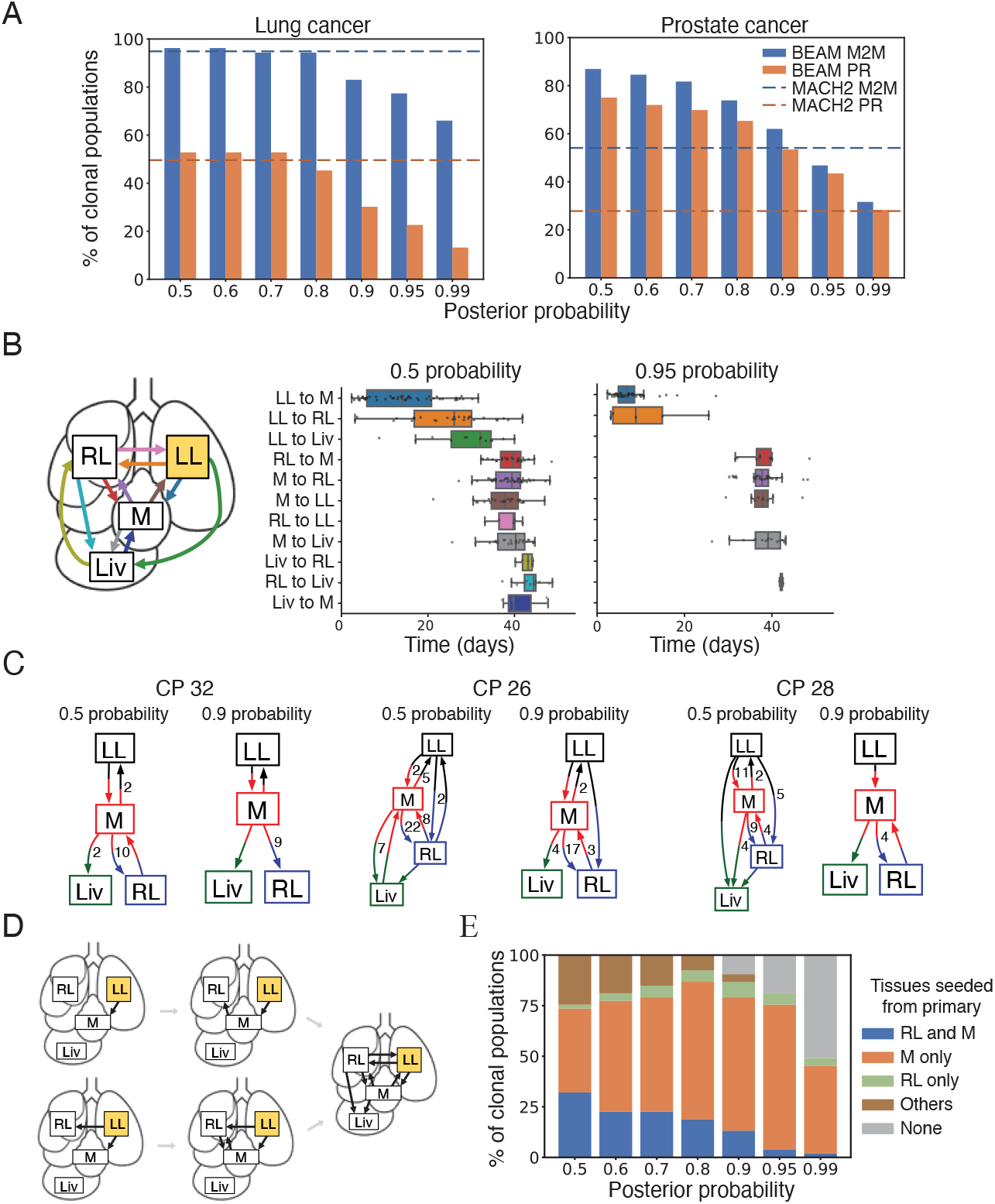
**A.** Fractions of clonal populations (CPs) for which metastasis-to-metastasis (M2M) or primary reseeding (PR) events were detected by BEAM at various thresholds of edgewise posterior probability for the lung [34] (left) and prostate [37] (right) datasets. For comparison, estimates from MACH2 [51] (averaged across thresholds due to threshold invariance) are shown as dashed lines. **B**. Times estimated by BEAM for each type of migration in the lung cancer data at 0.5 and 0.95 edgewise posterior probability thresholds. Boxplots represent distributions of expected migration times per edge type per CP. Each expected time represents an average over posterior samples of midpoint times for all corresponding lineage-tree branches. **C**. Example 0.5 and 0.9-threshold consensus graphs for lung cancer CP32, CP28 and CP26, illustrating the M-hub and direct LL→RL models. Numbers to the right of edges indicate counts >1 of corresponding branches in cell lineage trees, or equivalently, the multiplicity of the edges in the multigraph. **D**. Representative progression of M-hub and direct LL→RL models. **E**. Classification of all lung cancer CPs with increasing posterior probability threshold. Cases are classified first as LL→RL and LL→M (RL and M), then M only (all of which are LL→M), then RL only (all of which are LL→RL), and lastly based on whether or not LL seeded any other tissues (Others or None). RL = right lung, LL = left lung, M = mediastinum, and Liv = liver.

The prostate cancer data differed in several important ways from the lung cancer data. For various reasons—including the use of the SEMM with immunocompetent mice, a DNA readout, and a longer-duration experiment (up to 60 weeks) with more opportunity for drop-out of mutated cells—many fewer barcode mutations are captured in this system, making the phylogenetic inference problem considerably more challenging. In addition, a more diverse set of organs was sampled in this study, including the prostate, liver, lungs, bones, bladder, and lymph nodes, whereas the lung cancer data were dominated by three organs in close proximity (right and left lung, mediastinum). In part for these reasons, and likely also owing to some differences in their analysis pipeline, Serio et al. [37] reported quite low rates of M2M and PR events, detecting them in only ∼7% and ∼0.3% of CPs, respectively.

In our reanalysis, both BEAM and MACH2 detected substantially higher rates of M2M and PR events in the prostate cancer dataset (**Fig. 3A**; **Suppl. Fig. S7**). MACH2 found M2M events in ∼54% of CPs and PR events in ∼28% of CPs. At a posterior-probability threshold of 0.8, BEAM estimated even higher frequencies of ∼74% and ∼65%, respectively. As with the lung cancer data, however, BEAM’s estimates depended strongly on the choice of threshold, declining from ∼87% to ∼32% for M2M and from ∼75% to ∼28% for PR as the threshold increased, indicating that many of the M2M and PR events had weak support in the data. As discussed in the next section, these estimates are undoubtedly influenced by limits in the mutational information available in this dataset, but overall they suggest that the previous analysis may have been somewhat conservative in detecting secondary migration events.

One advantage of our sampling-based Bayesian inference approach is that it naturally allows for estimation of the times at which tissue-migration events occur, within the limits of a molecular-clock assumption (**Materials & Methods**). When analyzing the lung cancer dataset, we noticed that the primary left lung (LL) tissue was often predicted to seed the lymphatic mediastinum (M) and right lung (RL) tissues early in disease progression (**Fig. 3B**). In further examination of the inferred migration graphs, we found that early migrations tended to follow one of two distinct patterns: one in which M was the only site seeded from LL and subsequently acted as a hub for all other migrations; and another in which both M and RL were seeded directly from LL (**Fig. 3C**). Although the early dynamics differed between these patterns, both eventually resulted in widespread dissemination (**Fig. 3D**). The proportion of CPs in each pattern changed with the posterior probability threshold, but nearly all CPs fit into one of the two observed patterns (**Fig. 3E**). We also observed that metastasis to the liver (Liv) was most likely to occur via M, rather than from LL or RL (**Suppl. Tab. S1**). Overall, these patterns were broadly consistent with a previously reported principal components analysis [34], but BEAM was able to provide additional information about the timing of migration events, further revealing the patterns by which these tumors spread

### Bayesian hypothesis testing addresses higher-level questions about cancer metastasis

The previous section illustrates a recurring issue in the analysis of metastasis: while reconstruction methods generally produce full tissue-migration graphs, investigators are often interested in higher-level questions, such as the frequency of M2M or PR events, or whether or not the mediastinum acted as a migration hub. We extended BEAM to address questions of this kind by averaging over possible migration graphs and weighting them by their posterior probabilities, using the framework of Bayesian hypothesis testing. In this way, particular hypotheses of interest can be tested without relying on one or a few reconstructed migration graphs, each with high levels of uncertainty.

To illustrate the utility of this approach, we returned to questions that arose in our reanalysis of the lung and prostate cancer datasets. First, we sought to test whether or not the prostate cancer dataset supported the hypothesis of PR of the prostate from other metastatic sites, given the discrepancies between migration history inference methods and across BEAM posterior distributions. In this case, however, we had the additional challenge of sparse mutational data, owing to constraints of the prostate-cancer mouse model. Indeed, we observed that a majority of the prostate cancer CPs exhibited either no phylogenetically informative mutations or similar numbers to the lowest mutation-rate category in our simulations, where inference was particularly challenging (left of **Fig. 2C**). We found the lung cancer data to be more informative, but still on the low end of our simulated mutation-rate categories. We observed similar limitations in mutation content in published data sets for metastatic pancreatic [44] and lung [35] cancer (**Supp. Fig. S8**), suggesting that generating enough mutations to enable robust migration-graph inference remains a general challenge in the field (see **Discussion**).

We therefore defined an initial hypothesis test to distinguish CPs that were sufficiently informative to support inference of tissue-migration histories from ones that were not. For this test, we compared an alternative model with a fully parameterized GTR tissue-migration model to a null model in which tissue labels were randomly sampled in proportion to their relative frequencies at the leaves of the tree. This test essentially evaluates whether or not the tissue labels evolve in a Markovian manner along the branches of the tree—that is, whether or not each cell’s tissue label depends on the label of its parent and the branch length between them. If such a dependency exists, then there is at least some information in the data about the tissue-migration process, whereas if it does not exist, no such information is present. When we applied this test to our simulated data, we found that it was minimally restrictive, allowing all but a few data sets to pass at a threshold of log Bayes factor [lBf] > 1.1 (**Suppl. Fig. S9**). Similarly, the test identified ∼98% of the lung cancer CPs as informative about tissue migration. When we applied it to the prostate cancer data set, however, we found that only four CPs (∼1%) passed the test (**Fig. 4A; Suppl. Tab. S2**), indicating that the limited mutational content in this data set permits only a small minority of CPs to be informative about migration history. These results aligned with a simpler measure of mutual information derived from the full posterior distributions over lineage trees and migration graphs (**Suppl. Fig. S10**).

**Figure 4:**
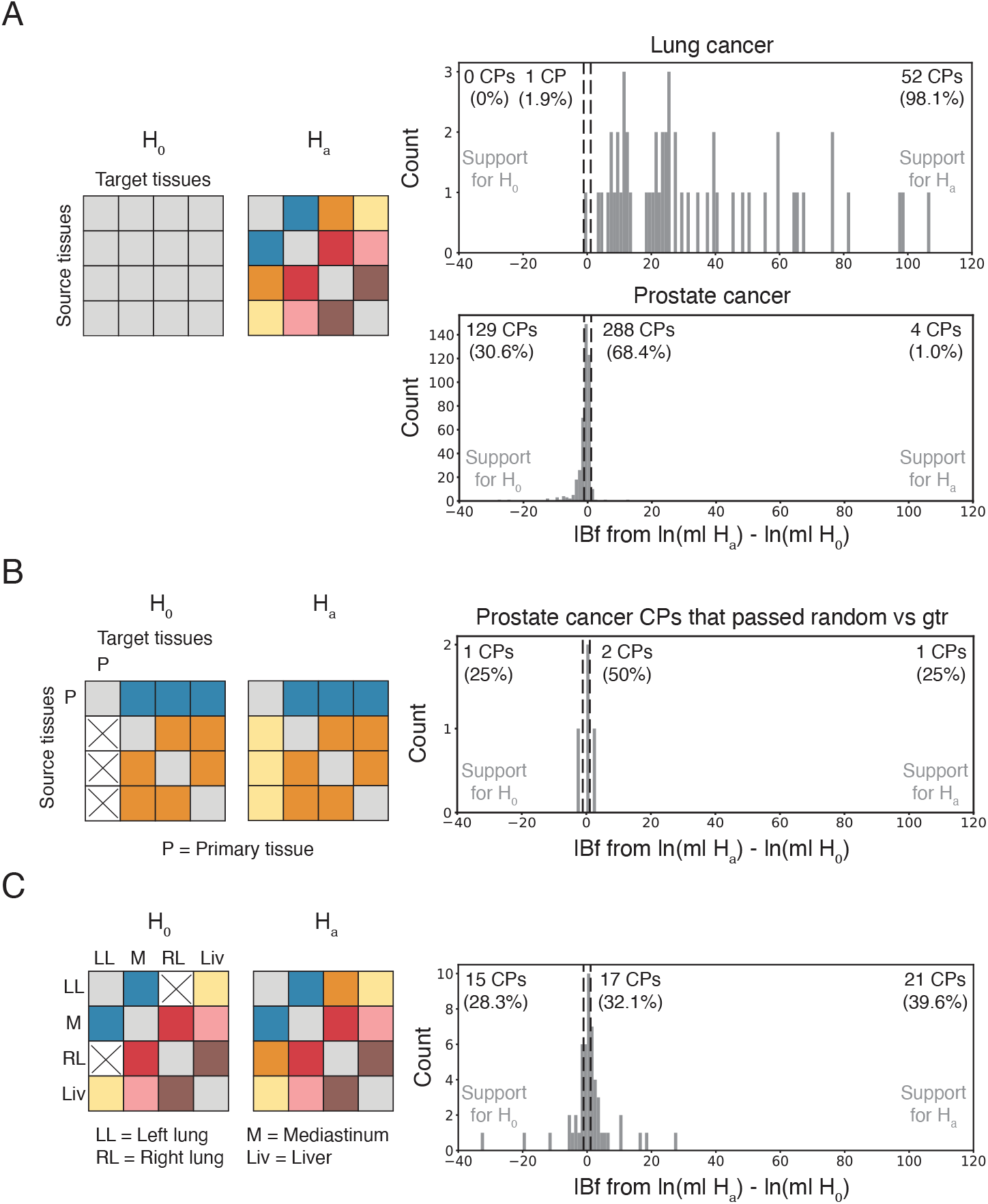
**A.** Hypothesis test of information about tissue migration based on a comparison of an alternative GTR model (right) against a random null model (left; see text). Results are shown for both the lung and prostate cancer datasets. **B**. Hypothesis test of an alternative model allowing for primary reseeding (right) against a null model prohibiting it (left). Results are shown for the prostate cancer CPs that passed the test in panel A. **C**. Hypothesis test comparing an alternative model allowing all types of tissue migration (full GTR model; right) against a null model prohibiting direct LL→RL seeding (left). CPs were each classified by log Bayes factor (lBf) into ones positively supporting the null model, the alternative model, or neither model. In all cases, thresholds of lBf *<* −1.1 and lBf > 1.1 were used (vertical dashed lines).

Based on this test, we focused our hypothesis-testing of PR on the four prostate cancer CPs that were identified as most informative about migration history. In this case, we tested an alternative model allowing for PR against a null model in which all PR events were forced to have rates of zero. We found positive support for PR in only one of the four CPs (**Fig. 4B**), which, interestingly, came from a different mouse than the single case of PR previously identified using MACHINA [37]. As validation, we performed the same hypothesis test on simulated datasets with and without reseeding events, and found that all simulated CPs predicted to have positive support for PR did indeed exhibit PR, while 75% of the simulated CPs predicted to have positive support for the null hypothesis did not exhibit PR (**Suppl. Fig. S11**). Among the 36 CPs without positive support in either direction, 18 of them (50%) did exhibit PR, suggesting that our test for PR is conservative. Thus, our Bayes-factor analysis does provide support that PR occurs in the prostate-cancer model, albeit in a small fraction of all CPs. Notably, the Bayesian hypothesis test is considerably more stringent than simply applying a posterior-probability threshold to the edges of the migration graph, as in the previous section.

For a second illustration, we designed a hypothesis test to distinguish between the two main patterns we observed in lung cancer. In particular, we tested an alternative hypothesis in which LL→RL events were allowed (corresponding to the blue bars in **Fig. 3E**) against a null hypothesis in which they were not, in which case we presume the data will primarily be explained by the M-hub model (orange bars in **Fig. 3E**). This test was implemented by comparing a fully parameterized model (representing the alternative hypothesis) against a model forced to have a rate of zero for LL→RL events (representing the null hypothesis). We found that the alternative hypothesis of direct LL→RL seeding obtained positive support (with lBf > 1.1) in ∼40% of eligible CPs, whereas the null hypothesis was positively supported (with lBf < −1.1) in ∼28% of CPs (**Fig. 4C**). In the remaining ∼32% of CPs, neither model was positively supported. This analysis further supports the finding that both the LL→RL and M-hub patterns are present in the data, without relying on particular migration graphs or posterior-probability thresholds. It additionally suggests that somewhat more CPs exhibit the LL→RL than the M-hub pathway.

## Discussion

In this article, we have introduced BEAM, a fully Bayesian framework that jointly models cell-lineage and tissue-migration histories to reconstruct the timing and routes of metastatic progression from single-cell lineage tracing data. To evaluate BEAM, we developed a data simulator that integrates an agent-based model of cancer metastasis with CRISPR-based lineage recording in DNA barcodes. We found that BEAM consistently recovered simulated migration histories with higher precision and recall than existing methods, and that the Bayesian approach was unique in allowing for flexible control over the confidence associated with each migration event. In real data for lung and prostate cancer, BEAM uncovered complex and highly connected migration graphs, even under stringent posterior-probability thresholds. In lung cancer, we identified two distinct patterns of metastatic progression driven by early divergence and found that liver metastases were preferentially seeded by secondary passage from a lymphatic tumor. In prostate cancer, BEAM highlighted diffuse posterior distributions over migration histories, which we traced to phylogenetic uncertainty resulting from limited mutational information. In both cases, BEAM effectively distinguished between signal and noise, preserving meaningful structure when data were informative while avoiding overfitting when they were not. We also showed that BEAM enabled direct hypothesis testing of features of the migration model, using it to assess the informativeness of lineage data, test for metastatic-to-primary reseeding events, and classify progression patterns.

Overall, BEAM makes three key contributions to methods for migration-history inference. First, it simultaneously addresses the intertwined problems of lineage-tree and migration-graph inference, enabling candidate lineage-trees to be evaluated in part by the likelihood of their induced migration graphs. Second, BEAM samples from a full posterior distribution over migration graphs, which supports tunable decision-making based on posterior-probability thresholds and avoids overconfidence in cases of weak data. Third, BEAM supports formal hypothesis testing by marginalizing over tree topologies and phylogenetic parameters, allowing researchers to rigorously assess questions about migration structure, data informativeness, and specific evolutionary events. Notably, BEAM is also modular by design, making it adaptable to new lineage-tracing technologies and experimental settings as they evolve. Together, these features make BEAM a uniquely powerful framework for extracting biologically meaningful insights from complex single-cell lineage tracing data.

In our re-analysis of publicly available lineage-tracing data sets, we observed that sparse information from mutations remains a persistent barrier to robust inference of tissue-migration histories (see also [58– 60]). This limitation was particularly acute with the prostate-cancer data set [37], which derived from an experimental system that was designed to maximize biological realism at some expense to barcode mutation rates. Nevertheless, we found that even more mutation-rich data sets for lung [34, 35] and pancreatic [44] cancer compared unfavorably with simulated data in the parameter regimes in which migration graphs could be reconstructed with high accuracy. Developing improved bar-coding techniques is a highly active area of research [61–65] (see [66] for a recent review), and we expect that the problem of sparse information will eventually fade in importance as experimental methods improve. At present, however, it is critical for investigators to ensure that reconstructed migration histories are well supported by the data before drawing strong biological conclusions from them. To our knowledge, the Bayes-factor-based test proposed in this article is the first formal statistical test for this purpose, and we anticipate that it and tests like it will be important in ensuring that migration-graph inference is well grounded in the available data.

An important distinction between BEAM and multi-criteria parsimony approaches such as MACHINA, MACH2, and Metient is that BEAM does not distinguish single-cell migration events from co-migrations of multiple cells, potentially from different parts of the lineage tree. El-Kebir et al. [45] introduced the idea of explicitly modeling co-migrations to address the problem that polyclonal events can cause the number of migrations to be over-estimated by standard methods. Their co-migration-aware formulation of the parsimony problem, however, is challenging to solve, and led them to employ a computationally intensive mixed integer linear programming approach using a commercial solver (Gurobi) (see also [50] and [51]). In principle, it would be possible to model co-migrations in our MCMC-based framework, but a good deal of additional complexity would be required. Instead, in this work we chose simply to model all migration events as occurring independently. Interestingly, even though co-migrations were frequent in our agent-based simulations, BEAM nevertheless performed well in recovering the true migration histories, suggesting that explicit modeling of co-migrations might not be necessary, provided a model is sufficiently flexible in other respects. Despite these encouraging results, it might still be of interest in future work to explore an extension of the Bayesian approach that separately models co-migrations. One interesting application of such a model would be to evaluate the degree to which the data supports the hypothesis of co-migration.

It is worth emphasizing that the question of modeling co-migrations is closely tied to an ongoing debate in the field surrounding polyclonal vs. monoclonal seeding of metastases. Early studies suggested metastases often arise from a single clone [67,68], whereas more recent studies have supported polyclonal origins, with distinct clones disseminating to secondary sites simultaneously or sequentially [28,31,45,69–73]. Nevertheless, polyclonal seeding does not appear to dominate in all cases [69,74]. For example, a large genomic study recently uncovered homogeneous, monoclonal metastatic sites for several cancer types [42]. Metastasis stage and therapy also influence clonal dynamics [28, 30, 32, 69, 74–77] with distant or post-therapy metastases often favoring monoclonal origins [30,76,78]. In cell-lineage tracing, both monoclonal and polyclonal lineage trees were observed within the same mouse, even when inferred using criteria that favored co-migrations [79]. Experimental sampling methods can also introduce bias when classifying monoclonal versus polyclonal origins [80]. Evidently, both monoclonal and polyclonal seeding occur across disease settings, calling for flexible modeling strategies that can accommodate both modes of tissue migration.

Another major source of computational complexity in methods that infer a migration graph by parsimony based on a given lineage tree is the presence in the starting tree of *polytomies*, or nodes with more than two children. Polytomies reflect uncertainty about the true structure of the tree and are common in trees estimated from lineage-tracing data, especially when mutations are sparse. To find parsimonious migration graphs, these methods have to consider the set of all possible refinements of polytomy-containing trees into binary trees, a set that grows exponentially with the number of polytomies. In practice, trees with many polytomies, such as those from the prostate cancer data set we analyzed, can lead to extremely long running times. BEAM circumvents this problem by directly exploring the space of tissue-labeled binary lineage trees and considering their likelihoods under both the barcode-mutation and tissue-migration models. As a result, it can avoid regions of the solution space where the lineage tree is compatible with the mutation data but the migration graph is unlikely, leading to some improvements in efficiency.

Despite these advantages, BEAM has similar limits in scalability to the other available migration-history inference methods. On one hand, BEAM benefits from simultaneous consideration of lineage trees and migration graphs, native handling of binary trees, a simple model of migration, and relatively fast likelihood calculations (see **Materials & Methods**). On the other, its reliance on exploration of the full space of tissue-labeled lineage trees by MCMC prohibits it from scaling to thousands of cells. BEAM’s precise scaling limits depended on several factors including tree size, mutation data quality, and the observed tissue label distribution, but we found that it could typically resolve lineage trees with up to 300–500 cells and was generally difficult to apply beyond that scale. In our hands, these limits were roughly comparable to those of the leading parsimony-based methods, despite major differences in the Bayesian and parsimony algorithmic strategies, suggesting that these are approximately the current limits of the field and new innovations will be needed for major improvements in scalability (see [79] for one such recent effort). In future work, we plan to explore the use of variational inference in place of MCMC to improve the scalability of BEAM (e.g., [81, 82]).

The technology of CRISPR/Cas9-based lineage tracing is only about a decade old and its use in the study of metastasis is even more recent. Computational methods for addressing these problems are still in their infancy and many opportunities remain for improvements. Nevertheless, BEAM represents an important step forward by introducing simultaneous Bayesian inference of lineage phylogenies and tissue migration graphs, as well as Bayesian hypothesis testing, to the field. Because uncertainty about the true structure of the lineage tree and migration graph tends to be high, the Bayesian approach is particularly powerful. We expect the ideas introduced here to open up new avenues for continued methods development, and to help enable a deeper understanding of the complex process of metastasis.

## Materials & Methods

### Modeling barcode mutation and tissue migration as conditionally independent processes

The barcode-mutation and tissue-migration processes were modeled as conditionally independent continuous-time Markov chains (CTMCs), given the phylogeny and branch lengths. Consider a single branch of the tree with length *b*, leading from a parent node *u* to a child node *v*. We represent the joint conditional probability of a barcode mutation state *B*_*v*_ and tissue label *T*_*v*_ at the child, given a barcode mutation state *B*_*u*_ and tissue label *T*_*u*_ at the parent, as a product of the two conditional probabilities,

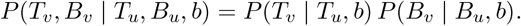

As a result, the likelihood of observed barcode data **B** and tissue labels **T** at the tips of a tree with topology 𝒯 and branch lengths **b** can be expressed as a product of phylogenetic likelihoods for the barcode and tissue labels, respectively,

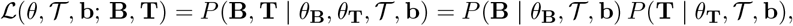

where *θ* = {*θ*_**T**_, *θ*_**B**_} such that *θ*_**B**_ contains parameters for the barcode mutation process and *θ*_**T**_ for the tissue migration process. Each CTMC is defined by a corresponding rate matrix and an overall strict clock rate, as detailed below.

The barcode mutation model, adapted from TiDeTree [53] and similar to LAML [19], was designed to describe an irreversible CRISPR-induced mutation process with the potential for silencing at individual sites. In particular, the model is defined by an infinitesimal generator *Q*_*B*_, such that:

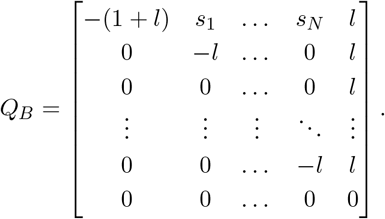

The first state represents the unmutated barcode, the last state represents a heritably silenced barcode, and the remaining *N* states represent unique indel (insertion or deletion) outcomes. Notice that all edits and silencing are assumed to be irreversible. Model parameters include the silencing rate *l* and relative indel rates **s**_*N*_, where *s*_*i*_ ∈ **s**_*N*_ is scaled such that 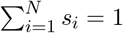. This scaling ensures that the expected editing rate is one and therefore that estimated branch lengths can be interpreted in units of expected indels per site. We omit the notion of a “scarring window” used in ref. [53] and allow mutations to occur at any time.

Tissue transitions were modeled with a separate infinitesimal generator *Q*_*T*_, whose state space corresponds to the set of available tissues. Each off-diagonal entry *q*_*ij*_ in *Q*_*T*_ represents the instantaneous migration rate from tissue *i* to tissue *j*. By default, we assume a general time reversible (GTR) parameterization, where each entry is given by *q*_*ij*_ = *r*_*ij*_*π*_*j*_, with symmetric exchangeability rates *r*_*ij*_ = *r*_*ji*_, and equilibrium tissue frequencies *π*_*j*_ such that Σ_*j*_ *π*_*j*_ = 1. The full rate matrix takes the form:

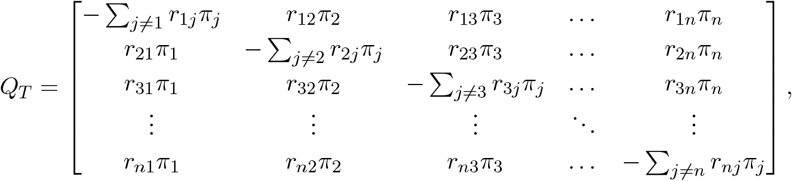

where the diagonal terms ensure that each row sums to zero. The full rate matrix is normalized such that, under the tissue equilibrium frequencies, the expected number of transitions per unit time is equal to one. BEAM explicitly parameterizes the equilibrium frequency of the first tissue, *π*_1_, which is assumed to be the primary source of the tumor, and sets the remaining frequencies to be equal:

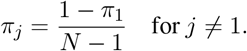

Likelihoods were computed using Felsenstein’s pruning algorithm [54]. In the case of the barcode mutation model, for a leaf node *i* with observed barcode state *B*_*i*_, the partial likelihood *L*_*i*_(*x*)—indicating the probability of the data beneath node *i* given that node *i* has state *x*—is initialized as:

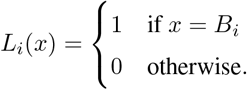

In the case of the tissue migration model, *L*_*i*_(*x*) is analogously set to 1 if, and only if, *x* is equal to the tissue label at leaf *i, T*_*i*_. In both cases, the recurrence relation for internal node *j* with children *k* and *l* is:

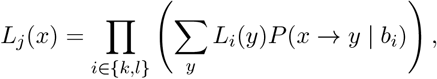

where *P* (*x* → *y* | *b*_*i*_) represents the conditional probability of state *y* at child node *i* ∈ {*k, l*} given state *x* at parent node *j* and branch length *b*_*i*_. These conditional probabilities are obtained, in the usual way, by computing the matrix exponential *P*_*B*_(*b*_*i*_) = exp(*Q*_*B*_*b*_*i*_) or *P*_*T*_ (*b*_*i*_) = exp(*Q*_*T*_ *b*_*i*_) [4].

At the root node *r*, the final likelihood is given by:

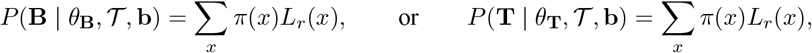

where *π*(*x*) is the equilibrium frequency of root state *x*. Because, in our setting, we typically can assume that the root has the unmutated state, we force *π*(*x*) to be equal to 1 for that state and zero otherwise. For the tissue migration process, we do the same for the primary tissue.

A general implementation of the pruning algorithm was used for the tissue-migration process, but for the barcode mutation process, optimizations were available due to irreversibility of the process. In particular, under this model, many ancestral states are not possible and can be excluded to improve efficiency [19]. Let *S* be the set of all observed indel states at the tips of the tree, including the unedited state (0) and missing data (−1), and let *S*_*j*_ be the set of observed descendant states for internal node *j*. The allowed ancestral states at node *j, A*_*j*_, are:

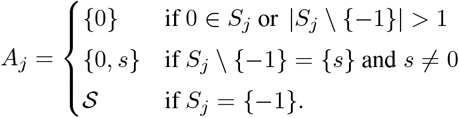

The summation in the recursive likelihood calculation can therefore be restricted to *y* ∈ *A*_*j*_, assuming that *L*_*j*_(*y*) = 0 for *y ∉ A*_*j*_, substantially improving runtime.

With these assumptions, implementation in BEAST 2 was straightforward. For each sampled or proposed tree topology 𝒯, branch lengths **b**, barcode mutation parameters *θ*_**B**_, and tissue-migration parameters *θ*_**T**_, we simply calculated the unnormalized posterior density by multiplying the prior densities and the phylogenetic likelihoods of the conditionally independent barcode data and tissue labels,

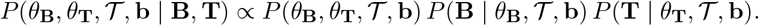

We relied on the existing functionality in BEAST 2 to explore the space of tree topologies and parameters. Convergence was assessed by checking that the effective sample size for all parameters exceeds 200, and by visually confirming that parameter traces had reached stationarity.

### BEAM parameters and priors

BEAM includes several free parameters that govern the tree topology, mutation dynamics, and tissue migration process. The phylogenetic tree topology is denoted by 𝒯, with associated branch lengths **b** = (*b*_1_, *b*_2_, …, *b*_*n*_). The prior over tree topologies and branch lengths is modeled using a birth-death process, with birth rate *λ* and death rate *µ*, which are treated as free parameters. The initial tree 𝒯_0_ can be provided directly in Newick format. In this study, we used LAML to infer a starting tree by approximate maximum likelihood. The barcode mutation model includes a silencing rate parameter *l*, while the relative rates of non-silencing edit outcomes, *s*_1_ … *s*_*N*_, are fixed and normalized based on the relative frequencies in the observed mutation matrix, as similarly done in [19, 53]. A strict molecular clock rate *ν*_*b*_ converts real-time branch lengths, which are directly operated on by MCMC proposals for a tree with fixed height from a specified experiment duration time, into number of substitutions before applying the rate matrix exponential to compute transition probabilities for the branch. Tissue migration dynamics are modeled with the relative migration rates between tissues denoted *r*_*ij*_, and a strict clock rate for tissue migration is represented by *ν*_*t*_. The equilibrium frequency of the primary tissue is denoted *π*_1_, with the frequencies of the remaining tissue types set to a uniform value to ensure that all frequencies sum to one.

The prior distributions assigned to these parameters were as follows:

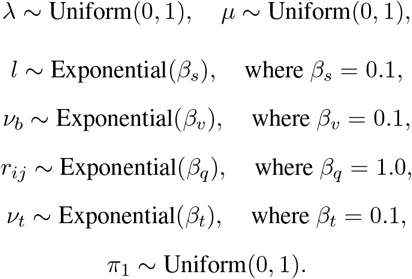

Initial values for parameters were selected based on empirical convergence behavior observed in pilot runs. We used minimally informative priors in this study, but BEAST 2 allows these prior distributions to be adjusted easily, as needed.

### Simulation model for cancer evolution and CRISPR lineage tracing

We developed an agent-based model of cancer cell dynamics built on three tools: the Tool for Tumor Progression [55], the MACHINA simulator [45], and the Cassiopeia barcode simulator [14]. MACHINA extended the original tumor progression model to include metastasis. Below, we briefly describe the key modeling components.

Cell birth and death follow a multiplicative fitness landscape with logistic constraints. The birth rate of cell *c* is

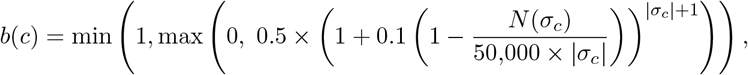

where *N* (*σ*_*c*_) is the number of cells in the same tissue with driver mutation set *σ*_*c*_. A cell divides if *r ∼* Uniform(0, 1) *< b*(*c*); otherwise, it dies. Upon division, one daughter retains the parent’s genotype, while the other mutates with probability *p* = 0.1. Mutations are drivers with probability *p*_*d*_ = (2 × 10^−7^)(|*σ*_*c*_| + 1), and passengers otherwise. Each generation, birth/death decisions are made for all cells, followed by metastasis decisions for each existing tissue. The probability of migration from tissue *t* is

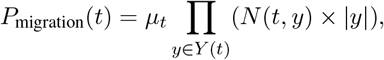

where *N* (*t, y*) is the number of cells in tissue *t* with driver mutation set *y*, and *µ*_*t*_ is the migration rate per cell per driver. Migration occurs from a tissue if *r ∼* Uniform(0, 1) *< P*_migration_(*t*) and the number of migrating cells is then drawn from Poisson(1) and migrated to a destination tissue chosen from a uniform transition matrix across ten tissues.

Simulations run for 250 generations, after which cells are downsampled. The lineage tree with branch lengths in cell divisions is then used to simulate CRISPR barcodes at the tips using Cassiopeia, assuming a uniform mutation rate across sites, default heritable silencing rate of 0.0001, and default stochastic silencing rate of 0.01. The output includes a cell-by-barcode matrix, tissue labels, and ground-truth phylogeny and migration history.

### Simulated data

To generate simulated datasets under ideal conditions, we ran 100 simulations with a mutation rate of 0.0025 mutations per barcode site per cell division and a migration rate of 1 × 10^−6^ per cell. These simulations produced a ∼50% saturated character matrix and a migration graph of intermediate complexity (see representative examples in **Suppl. Fig. S2**). To simulate data with variable complexity of barcode mutation and tissue migration, we simulated 20 datasets for each combination of barcode mutation rates {0.0005, 0.001, 0.0025, 0.005, 0.01} per barcode site per cell division and tissue migration rates {1 × 10^−7^, 1 × 10^−6^, 1 × 10^−5^, 1 × 10^−4^} per cell (representatives shown in **Fig. 2C**). We simulated trees with 50 cells after downsampling. Simulations in which no migrations occurred were discarded and replaced with new ones until the desired number was reached. All simulations were processed with all migration history inference methods with the exception of three simulations with MACHINA due to long runtimes (two simulations for the 0.0005 mutation rate and 1 × 10^−7^ migration rate category and one simulation for the 0.0005 mutation rate and 1 × 10^−4^ migration rate category).

### Alternative methods for benchmarking

For consistency and efficiency in benchmarking, we developed a pipeline to process the same simulated datasets by several methods in parallel. Because all methods except BEAM required a fixed lineage phylogeny as input, we selected a single phylogeny inference method for preprocessing and, for each simulated mutation matrix, we applied this method once and used the resulting tree for all downstream steps. Based on published benchmarks and our preference for a probabilistic reconstruction method, we chose the LAML method [19] for phylogeny inference. Notably, LAML uses an approximate maximum-likelihood method to estimate both a topology and branch lengths under nearly the same mutation model as the one assumed by BEAM. For initialization of LAML, we used the Cassiopeia-Greedy algorithm [14] to generate an approximate tree topology without branch lengths from the mutation matrix.

Once this cell-lineage tree was obtained, we applied the following methods for migration-graph inference (listed in order of increasing sophistication):

#### Random

Tissue labels at the tips of the input phylogeny were used to randomly assign tissue states to internal nodes, ignoring the tree structure. This approach served as a baseline for migration-graph accuracy under random ancestral state assignments and provided an approximate lower bound for the performance of other methods.

#### Consensus

For each internal node, tissue labels for all tips in its sub-tree were collected and the most common tissue label among those tips was assigned to the node. This was the simplest possible use of the tree structure for tissue labeling.

#### Parsimony

We applied our own implementation of the Fitch-Hartigan algorithm [56, 57] for the “small parsimony problem” of discrete ancestral state reconstruction. This method minimized the number of migration events on an input cell lineage tree, without refining that tree to resolve polytomies or considering other parsimony criteria.

#### MACHINA [45]

We applied MACHINA v1.0 in Parsimonious Migration History with Tree Refinement (PMH-TR) mode to both refine a given clone tree into a binary tree, resolving polytomies, and label its internal nodes with tissue assignments to infer a migration history. The MACHINA objective is to minimize, in order: the number of migration events (directed edges in the migration graph), the number of co-migrations (unique edges in the migration graph, so that a single directed edge and a directed multi-edge each only contribute a count of one), and the number of seeding sites (distinct tissues serving as sources of migration) using mixed integer linear programming under a parsimony criterion. We used MACHINA in unrestricted migration mode to allow the migration graph to take any topology rather than being restricted to predefined patterns. MACHINA returned a single most parsimonious labeling.

#### Metient [50]

We applied Metient in a similar way to MACHINA. Metient tries to minimize the same three criteria of migrations, co-migrations, and seeding sites while resolving polytomies in the tree, similar to MACHINA, but it allows flexibility in how these criteria are weighted. We ran Metient-evaluate using the default parsimony weights (0.48 for migrations, 0.30 for co-migrations, and 0.22 for seeding sites), which were optimized in the original method across across several cohorts of cancer patients. We also tested Metient-calibrate on our simulated data, which found different optimal weights but led to worse performance, so we report results using the default weights from Metient-evaluate. Tissue locations for tips were encoded in the input metadata as present and nodes were marked absent from all tissue locations. For precision and recall based performance metrics, the top migration graph solution was selected based on the minimum of the solutions’ loss scores, as in [50].

#### MACH2 [51]

Finally, we applied MACH2, an extension of MACHINA which solves the same PMH-TR problem but enumerates the full set of equally parsimonious migration histories rather than returning a single solution. All solutions are derived from the same integer linear programming formulation used in MACHINA, but MACH2 leverages combinatorial properties of the solution space to exhaustively enumerate all optimal solutions. We ran MACH2 in the default mode for v1.0.1 and the output solutions were all weighted equally (as in [51]) to construct a consensus migration graph summarizing the full solution set.

### Precision, recall, and F1 score calculations

To evaluate migration graph inference, we assessed the precision and recall of inferred edges in terms of true positive (TP), false positive (FP), and false negative (FN) edges in the graph. Importantly, both the simulated and inferred migration graphs are actually *directed multigraphs*, with the possibility of multiple edges between each pair of nodes. Therefore, motivated by ref. [51], we computed precision and recall using *counts* of each type of edge per graph rather than by considering their presence/absence. This strategy penalizes cases where the types of migration events are correctly inferred but the numbers of events are not—a kind of error that is particularly relevant when evaluating the clonality of seeding.

Given a true migration graph *G* and an inferred migration graph *G*^∗^ let *E*_*ij*_ and 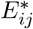 represent the numbers of directed edges in *G* and *G*^∗^, respectively, that run from tissue *i* to *j* where *i≠ j*. The counts TP, FP, and FN were derived from these edge counts as follows:

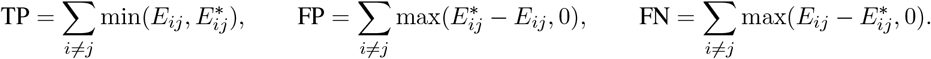

The precision (Pr), recall (Re), and F1 score were then calculated by the standard formulas:

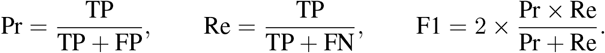

For precision-recall curves from MCMC samples, we first converted each sample *s* of a tissue-labeled tree into a migration multigraph, with an integer count 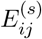 for each type of edge (*i, j*). We then let *f*_*ij*_(*c*) be the fraction of such sampled multigraphs that have *c* or more counts of edge (*i, j*), that is, the fraction with 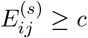. Because it is derived from samples from the approximate posterior distribution, this fraction *f*_*ij*_(*c*) can be interpreted as an estimate of the posterior probability that the inferred edge count 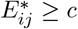. We therefore subjected this value to a varying posterior-probability threshold, *p*, such that 0 ≤ *p* ≤ 1. For each choice of *p*, we estimated 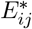 as the maximum number of edges *c* such that *f*_*ij*_(*c*) ≥ *p*. We then evaluated the counts TP, FP, and FN using that value of 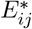. For the F1 score calculations, we fixed the threshold *p* at 0.5.

We used the same strategy for MACH2, treating each reported solution as if it were a draw from a posterior distribution (so that they were equally weighted). For Metient, however, we followed the authors in using the single solution that minimized their loss function [50].

We followed ref. [83] in using precision-recall curves rather than receiver operating characteristic (ROC) curves, which are problematic in the multigraph setting.

### Calculating excess migrations relative to parsimony

To calculate the excess migrations predicted by BEAM, we began with a sample from the joint posterior distribution over tissue-labeled trees and model parameters given the data. For each sampled tree, the number of BEAM-inferred migration events was obtained in the usual way, by counting branches in the tree for which the parent and child tissue labels disagreed. The tissue labels were then removed from the internal nodes of the tree and our implementation of the Fitch-Hartigan parsimony algorithm was applied to obtain an estimate of the minimum possible number of migration events given the leaf labels only. This minimum possible count was subtracted from the BEAM estimate to obtain an estimate of the excess migrations per sample. Finally, an estimate of the posterior expected excess migration count was obtained by averaging these per-sample excess counts across all samples.

### Application to lung, prostate, and pancreatic cancer datasets

As detailed in ref. [34], the lung cancer data was derived from orthotopically xenografted KRAS-mutant A549 lung adenocarcinoma cells, engineered with CRISPR barcodes and surgically implanted into the left lungs of four mice. As in the published analysis, we focused on data for one mouse (labeled “5k”) with 100 clonal populations (CPs), which were tracked over 54 days. We began with the provided allele table and metadata for mouse 5k. We excluded the 17 CPs excluded in the original paper (5, 16, 18, 25, 33, 38, 39, 41, 50, 53, 65, 69, 75, 81, 87, 88, 93), 9 CPs with only one observed tissue label (22, 29, 46, 48, 49, 78, 85, 94, 96), and 21 large CPs (1–4, 6–15, 17, 19–23, 27, 31) that were computationally prohibitive to analyze, and analyzed the remaining 53 CPs with BEAM. To infer tissue migration graphs, we used the coarse-grained tissue annotations, specifically the metadata labels LL, M, RL, and Liv, which were mapped to each cell barcode.

The prostate cancer data was obtained from a somatically engineered mouse with knockout of PTEN/P53 [84] and the introduction of CRISPR barcodes. Cancer developed in 10 mice over an average of ∼380 days, producing a variable number of CPs per mouse. The prostate cancer datasets for all mice were re-processed using the EvoTraceR pipeline (v1.0.4), following the authors’ original analysis protocol, resulting in 522 CPs. We excluded 2 CPs (MMUS1875 CP1 and MMUS1466 CP1) that were computationally prohibitive to analyze and we excluded 99 CPs with only one observed tissue label (MMUS1492 CP4, 6, 7, 9, and 13; MMUS1588 CP10, 13, 19, 21; MMUS1469 CP67, 77, 84, 93, 96, 97, 100, 103, 114–116, and 123; MMUS1457 CP7, 11–14, 18, 19, 24–26, 28–30, 33, 34, 36, 38, 41, and 43; MMUS1875 CP9, 12, 16, 21, 22; MMUS1495 CP86, 89, 95–97, 104, 108, 109, 113, 115; MMUS1466 CP6, 15–17, 20, 22, 24, 26, 27; MMUS1544 CP36, 37, 40, 53, 55, 60, 62, 68, 70, 79, 82–86; MMUS1874 CP7, 18, 20, 22, 24, 26, 27, 28, 32–34; MMUS1467 CP12, 15, 17, 21, 26, 27, 29, 34, 35). This left us with 421 CPs to analyze across the 10 mice with BEAM. With MACH2 analyses, we also had to exclude 3 more CPs (MMUS1469 CP1 and 3; MMUS1495 CP1) due to long runtimes.

For both datasets, these preprocessing steps produced mutation matrices and associated tissue labels for each CP. We then inferred a baseline tree for each CP using the Cassiopeia-Greedy algorithm [14] and supplied this tree to LAML [19] for tree and branch-length estimation by maximum likelihood. These LAML-estimated lineage trees were later used both as the starting trees for BEAM and as the fixed input trees for all other migration-graph inference methods.

We incorporated two additional reference datasets for comparison of CRISPR barcode mutational content in metastatic CPs. From a published pancreatic cancer dataset [44], we retained 41 metastatic CPs after filtering out single-tissue clones, using the provided mutation matrices directly. From an additional lung cancer dataset [35], we retained 12 metastatic CPs after similar single-tissue clone filtering, deriving mutation matrices from the provided overall allele table.

### Previously reported levels of M2M and PR for real data

For the lung cancer dataset, we used the numbers of CPs reported to exhibit metastasis-to-metastasis (M2M) seeding and primary reseeding (PR) under the labels “metastatic cascade” and “reseeding” in Fig. 7 of ref. [34] across all CPs that they analyzed. For the prostate cancer dataset, however, the published analysis reported fractions of edges in full lineage trees within or across CPs [37], which did not directly correspond to our metrics. Therefore, we obtained the MACHINA migration-graph output files from the authors and recalculated the percentage of CPs showing evidence of M2M or PR events across all CPs in their study. Specifically, a CP was labeled as M2M if it had any edge excluding the primary tissue, and as PR if it had an edge with a non-primary source tissue and the primary tissue as the target tissue.

### Hypothesis testing with Bayes factors

Bayes factors are a flexible and powerful means for model comparison and hypothesis testing in a Bayesian setting, but they require evaluation of the marginal likelihood of the data, integrating over parameters and any other latent variables. In our case, the marginal likelihood given the model *M* has the formidable form:

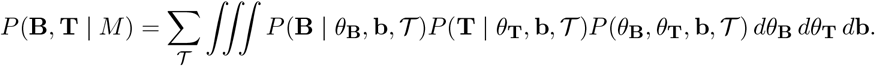

This expression—a version of what is sometimes called the “Felsenstein equation” [85, 86]—is both a sum over all possible tree topologies and an integral over all possible combinations of branch lengths and other parameters, making it highly intractable.

Nested sampling, however, offers an effective approach for approximating this integral by MCMC, and a convenient package for nested sampling is already available for BEAST 2 [87]. This approach works by mapping the parameter space to a prior mass using a cumulative density function [88], thereby transforming the high-dimensional integral into a one-dimensional integral over prior mass. The integral is approximated by sequentially sampling parameter sets of increasing likelihood and computing a weighted sum of these likelihoods, where the weights correspond to the changes in prior mass between samples.

Once marginal likelihoods are available for two models *M*_1_ and *M*_2_, a Bayes factor can easily be computed as a ratio of their marginal likelihoods. In the log space used by BEAST 2,

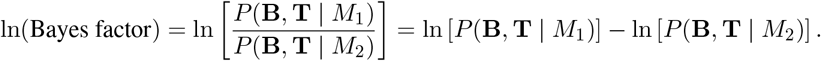

We simply interpreted values of ln(Bayes factor) *< −*1.1 as indicating positive support for *M*_2_, values of ln(Bayes factor) > 1.1 as indicating positive support for *M*_1_ and values between −1.1 and 1.1 (inclusive) as not positively supporting either model (see [89]).

In our case, we began by running nested sampling with 10,000 MCMC iterations per sub-chain and a single active particle. We then increased the number of active particles as much as computationally feasible and following the recommendations of the Taming the BEAST tutorial [90].

### Calculating mutual information of tissue transitions

For a simple, model-free evaluation of the information in the data relevant to migration history in the posterior distribution, we computed the mutual information between the sampled tissue assignments at parent and child nodes across all branches of the tree. First, a collection of trees sampled by BEAM was traversed to fill an *n × n* matrix of counts *C*, where *n* is the number of distinct tissue labels, and *C*_*ij*_ represented the total number of inferred transitions from tissue *i* to tissue *j* across branches. The count matrix was then normalized to obtain a matrix of joint probabilities of pairs of tissues, 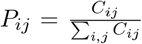, as well as vectors of marginal probabilities, *P*_*i*._ = Σ_*j*_ *P*_*ij*_ and *P*_.*j*_ = Σ_*i*_ *P*_*ij*_. The mutual information between the source tissue *X* and target tissue *Y* was then calculated as:

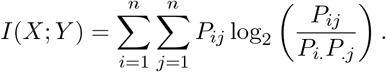

To normalize for differences in entropy between datasets, the mutual information was scaled by the average entropy of the marginals:

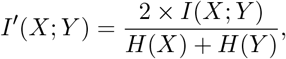

where *H*(*X*) = −Σ_*i*_ *P*_*i*._ log_2_(*P*_*i*._) and *H*(*Y*) = −Σ_*j*_ *P*_.*j*_ log_2_(*P*_.*j*_).

Since BEAM models tissue transitions as a Markov process, where ancestral tissue labels are more likely to persist within the same tissue along the cell lineage tree rather than to result from frequent migration between tissues, we expected a well-fit model to produce a high normalized mutual information score.

### Evaluating the PR test on simulated data

All ground-truth PR simulations from the variable rates simulated dataset in **Fig. 2C** were collected, excluding any with a mutation rate of 0.0005 or a migration rate of 1 × 10^−4^ due to poor performance in prior evaluations or a migration rate of 1 × 10^−7^ due to a lack of ground truth migration graphs with PR. This resulted in 39 valid PR simulations. All non-PR ground truth simulations were then gathered, excluding those with migration rate of 1 × 10^−4^, migration rate of 1 × 10^−7^, or mutation rate of 0.0005 and 39 of them were randomly sampled to match the number of PR simulations for class balance. We then applied the PR hypothesis testing procedure to all 78 simulations. Classification was based on the Bayes factor, where simulations with ln(Bayes factor) > 1.1 were classified as PR and ln(Bayes factor) *< −*1.1 as non-PR. This classification scheme ignores simulations classified as not supporting either model with −1.1 ≤ ln(Bayes factor) ≤ 1.1, so we provide results for all three outcome categories.

## Software availability

BEAM was implemented as a new set of Java classes for use within BEAST 2 [52]. The BEAM package is freely available as open source software under GNU General Public License v3.0 at https://github.com/StephenStaklinski/beam. A supplemental python package for working with BEAM output and running data simulations is available under the MIT License at https://github.com/StephenStaklinski/beam_sup.

## Data availability

The lung and prostate cancer data reanalyzed in this study were found in the original reports [34, 37]. All analysis pipelines were made publicly available at https://github.com/StephenStaklinski/bayesian_phylogenetic_metastasis.

## Acknowledgments

We thank David M. McCandlish, Bruce Stillman, and Hannah V. Meyer for their valuable feedback during the development of BEAM. We are also grateful to members of the Siepel lab for their helpful discussions. We thank Mrinmoy S. Roddur for assistance with MACH2 and Divya Koyyalagunta for guidance on using Metient.

## Supplementary Figures

**Figure S1:**
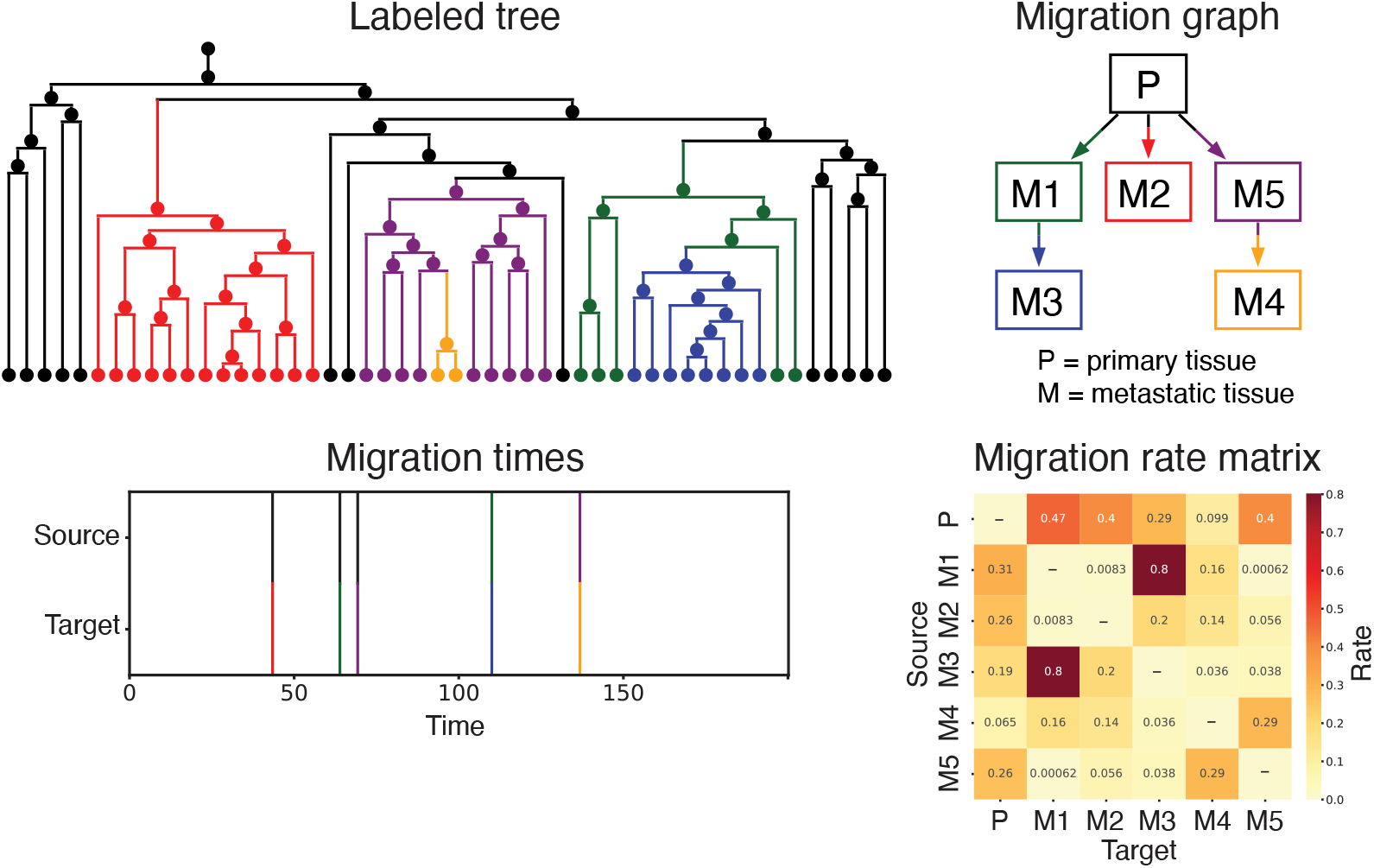
An example of a sample from BEAM’s posterior distribution, including a labeled tree (top left), the corresponding migration graph (top right) and derived migration times (bottom left). The migration rate parameters (bottom right) are also sampled from the posterior distribution.

**Figure S2:**
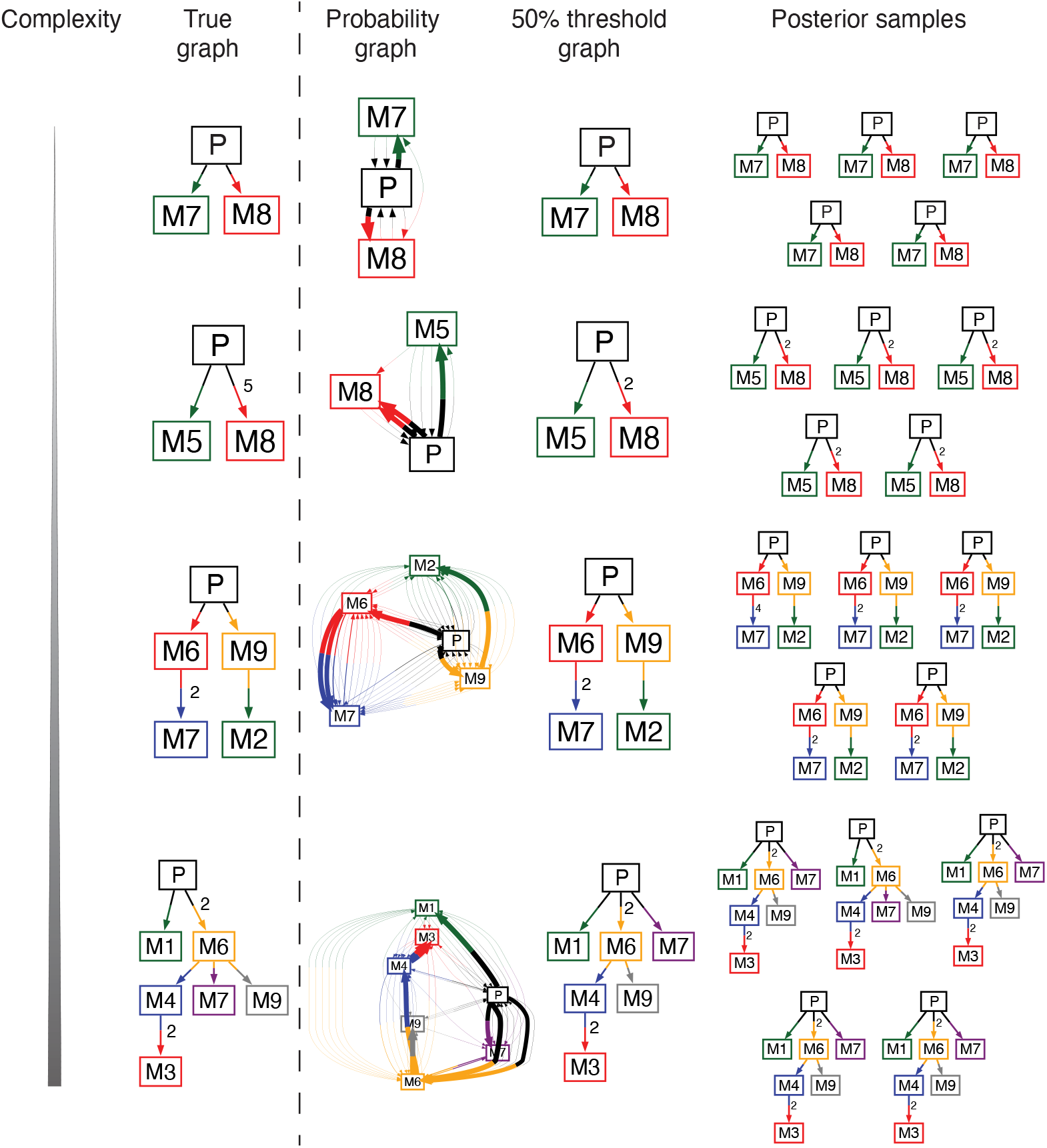
Representative graphs sampled from BEAM’s posterior distribution for simulated datasets of increasing complexity in the favorable parameter regime shown in **Fig. 2A**. Shown for each case are the true migration graph, the probability-weighted edge graph, the 0.5-threshold consensus graph, and individual samples from the posterior distribution. Numbers next to edges indicate multi-edge counts. Edges without a number represent single migration events. The P label represents the primary tissue and labels beginning with M are metastatic tissues in the simulations.

**Figure S3:**
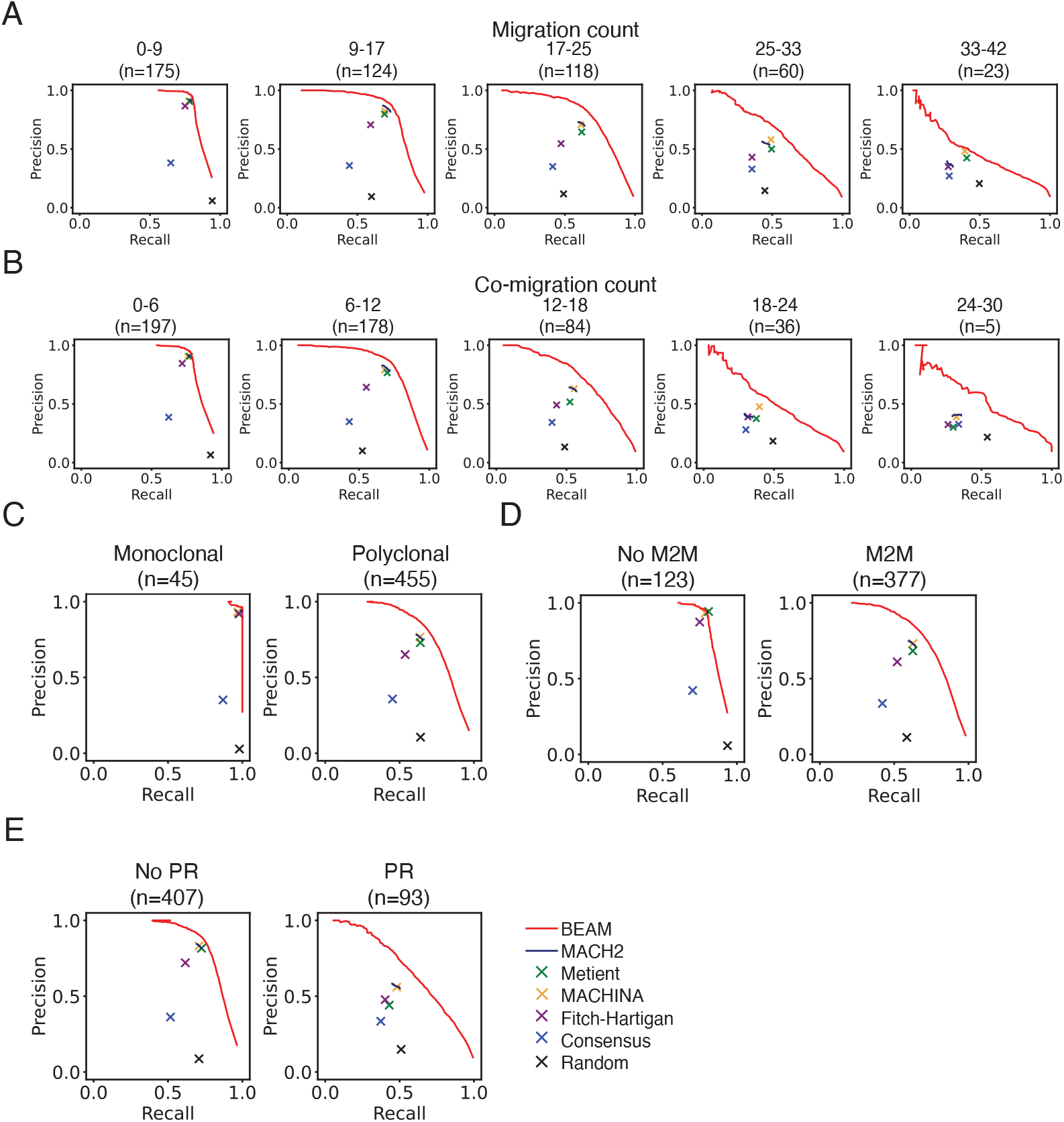
Precision-recall curves for all simulated datasets (500 total simulated data from the combination of **Fig. 2A** and **Fig. 2B-C**) stratified by: **A**. number of migration events (directed edges in the graph), **B**. number of co-migration events (unique edges in the graph, so that a single directed edge and a directed multi-edge each only contribute a count of one as in [45,50,51]), **C**. clonality (whether any multi-edges exist or not in the graph for polyclonal and monoclonal respectively), **D**. presence of metastasis-to-metastasis (M2M) seeding, and **E**. presence of primary reseeding (PR).

**Figure S4:**
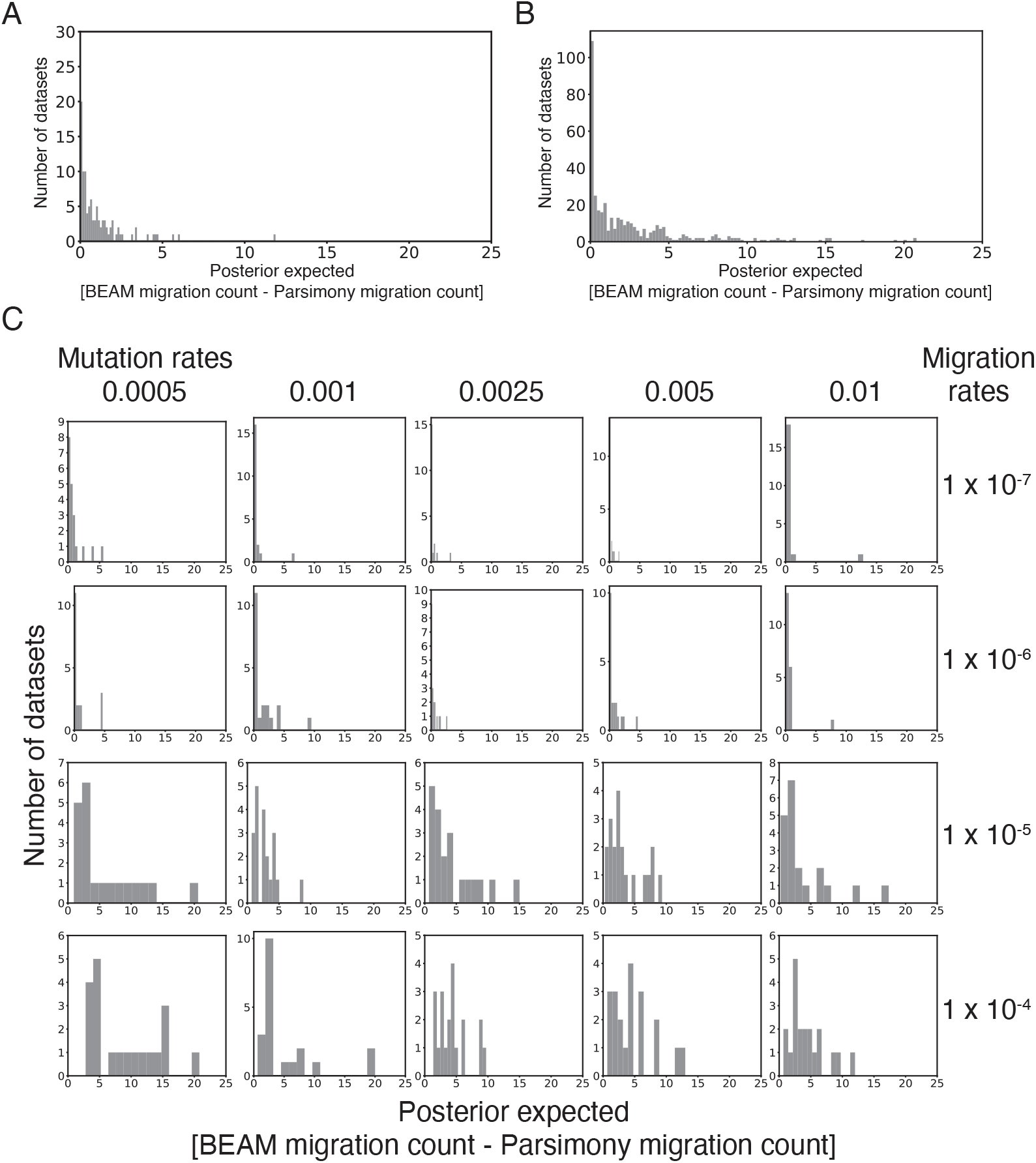
Histogram of the excess migrations predicted by BEAM relative to the Fitch-Hartigan parsimony solution for the same trees (see text) in: **A**. the favorable parameter regime shown in **Fig. 2A**; **B**. the variable parameter regime shown in **Fig. 2C**; and **C**. the variable parameter regime stratified by mutation and migration rate.

**Figure S5:**
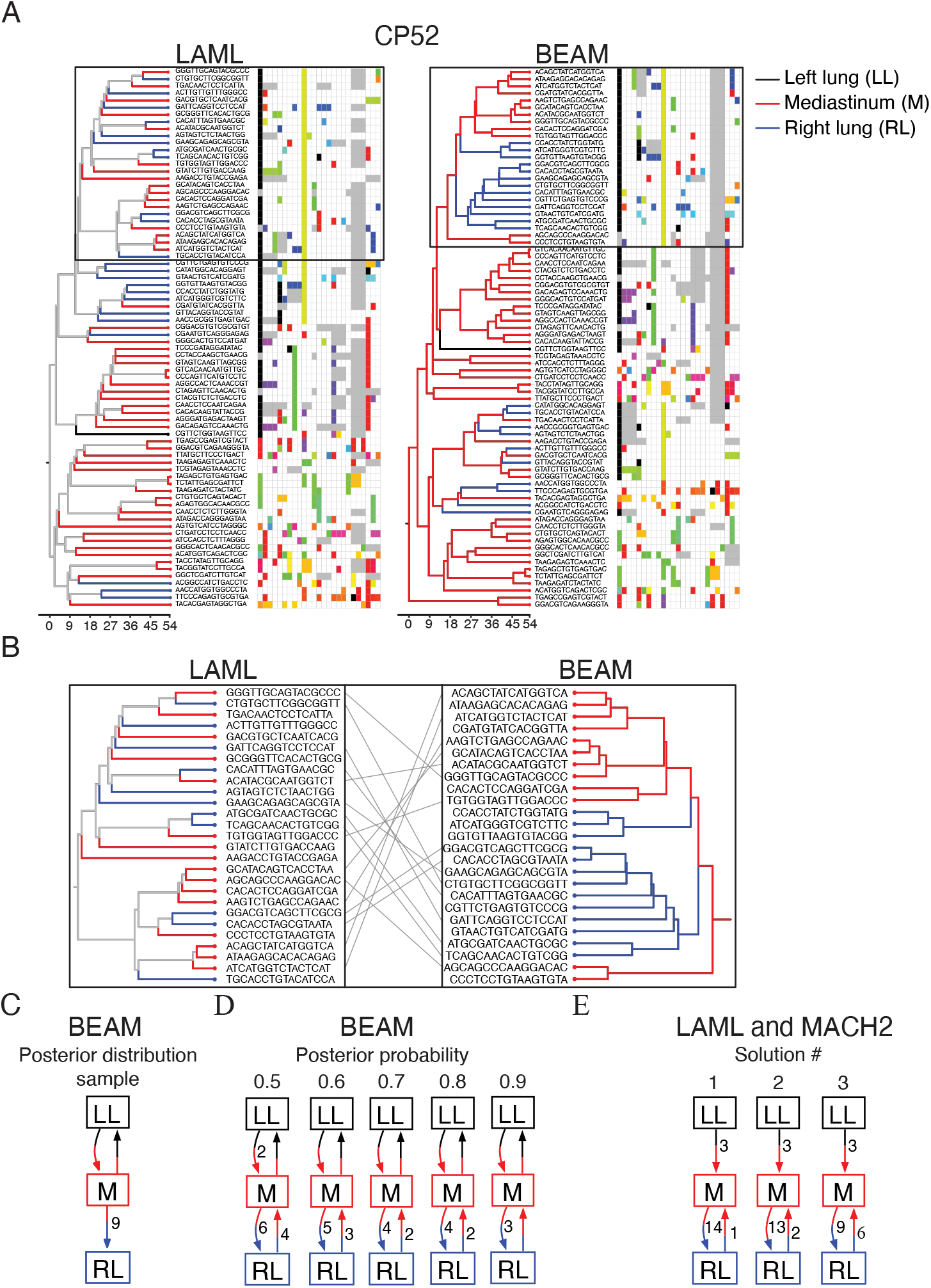
**A**. Lung cancer lineage trees for CP52 as inferred by LAML (left) and BEAM (right). LAML inferred only the tree topology, so internal nodes are gray, while BEAM resolved internal nodes by tissue. The matrix next to each tree shows barcode mutations per cell with white as unedited, gray as missing, and each color representing a mutation (with colors repeating if there are too many unique mutations). **B**. Zoomed-in view of a clade with topological differences between LAML and BEAM. Gray lines map tips present for both methods. **C**. Migration graph for the BEAM lineage tree in **A. D**. BEAM consensus migration graphs at increasing edge-retention thresholds. **E**. MACH2 migration graph resolved from the

**Figure S6:**
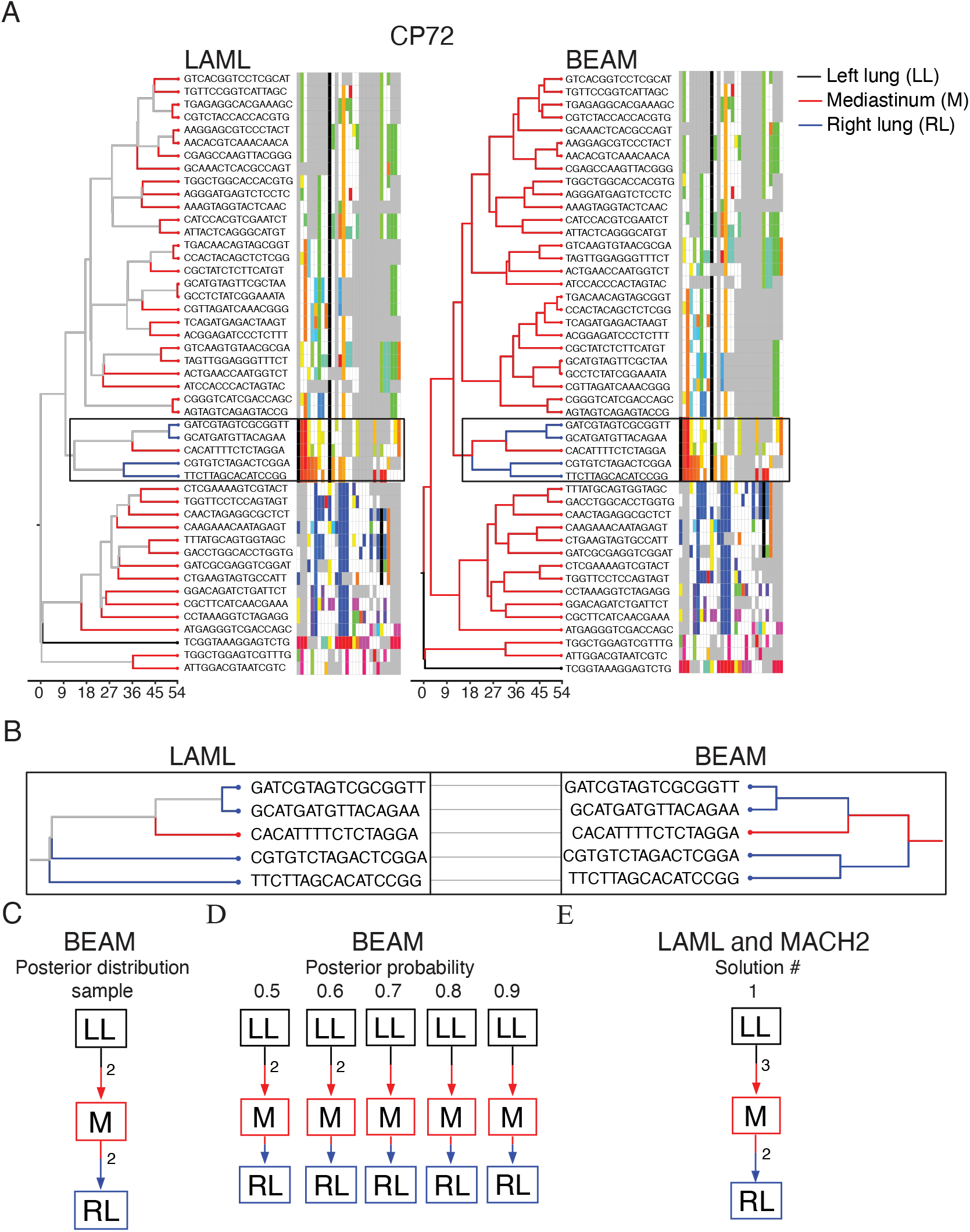
Same as **Suppl. Fig. 5** but for lung cancer CP72.

**Figure S7:**
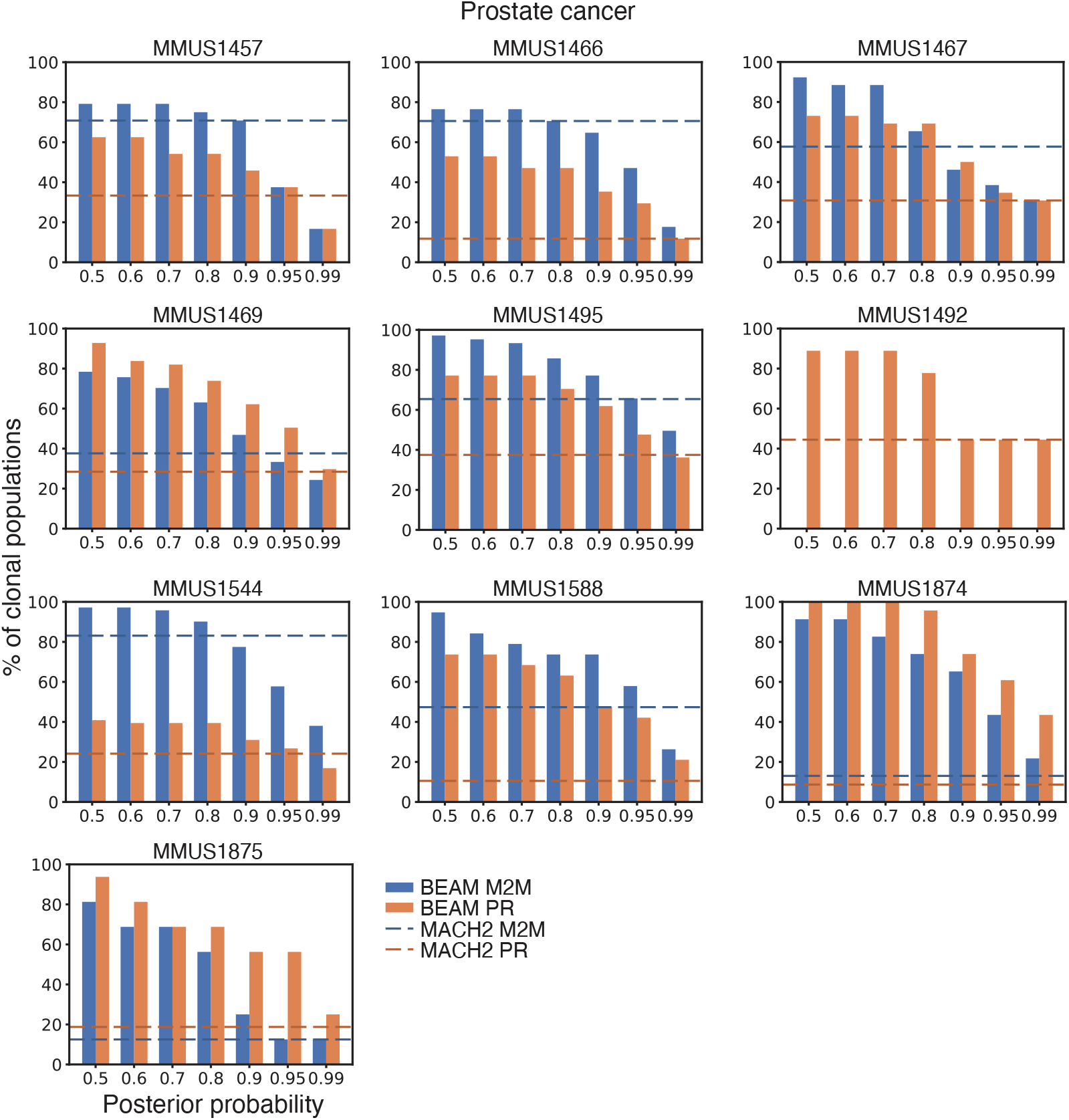
Same as **Fig. 3A** for the prostate cancer data plotted per mouse.

**Figure S8:**
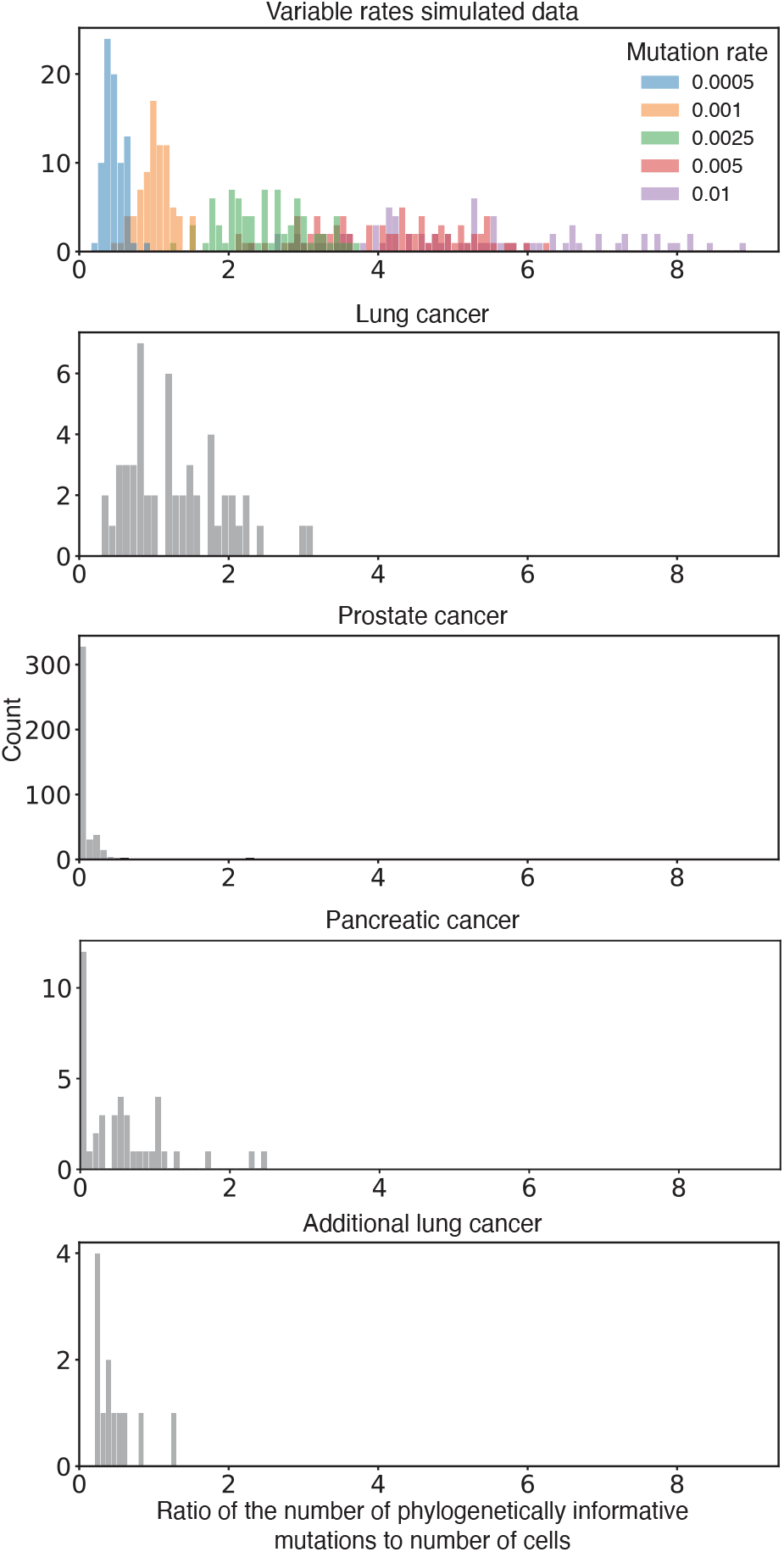
The ratio of phylogenetically informative mutations to cell number (tips in the tree) was computed for each clonal population in the real and variable-rate simulated datasets. “Lung cancer” and “Prostate cancer” correspond to the main datasets analyzed, while “Pancreatic cancer” and “Additional lung cancer” are additional external CRISPR barcode datasets [35, 44]. At each barcode site (column) in the mutation matrix, unique informative mutations were defined as the set of those mutations present in two or more cells (rows), but not all cells. Counts were summed across barcode sites and divided by the number of cells in the clonal population. Bars in the variable-rate simulated dataset are colored by simulated mutation rate for reference.

**Figure S9:**
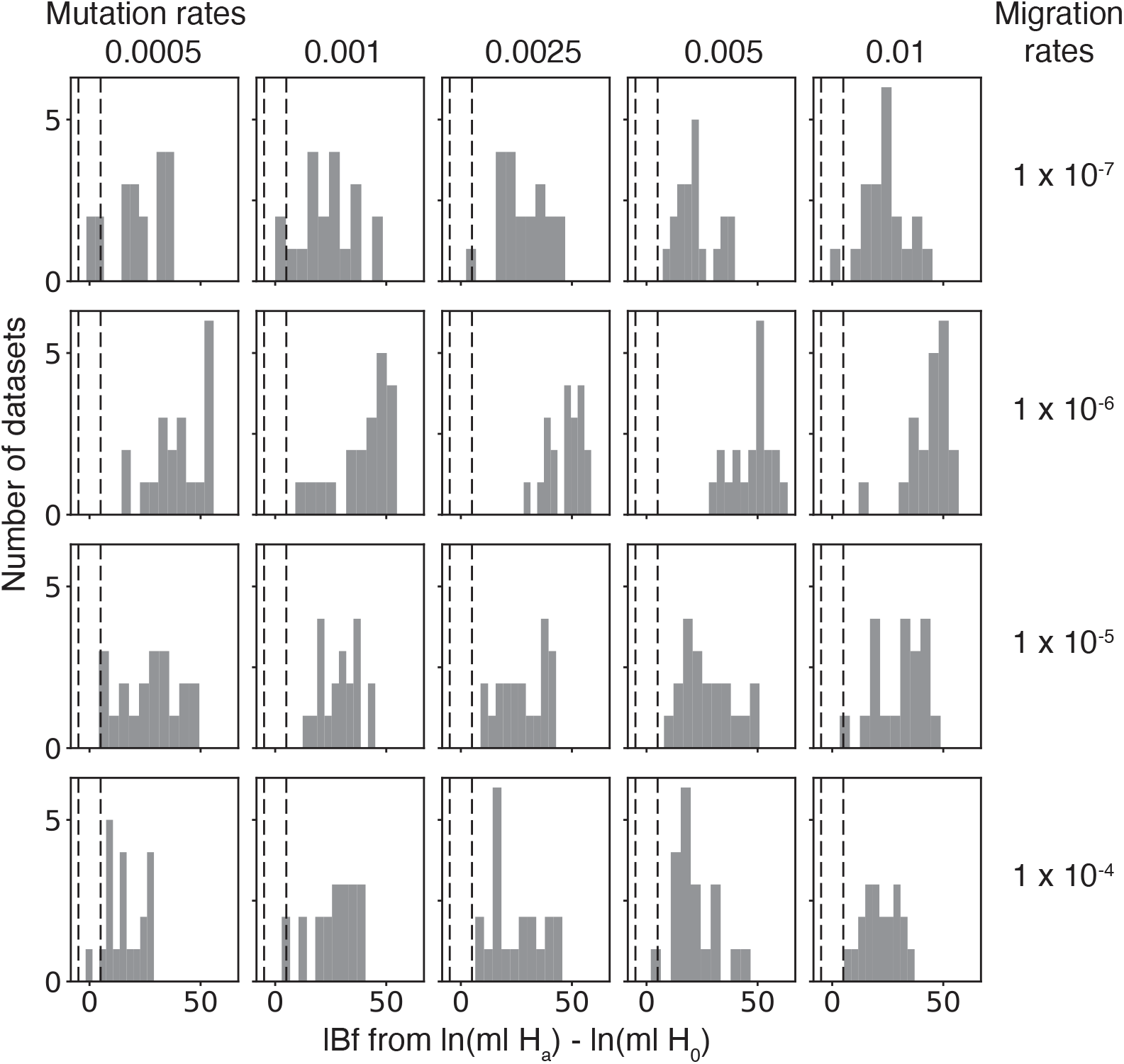
Application of the random vs. GTR information content hypothesis test in **Fig. 4A** to the variable rates simulated data from **Fig. 2C**. The reported values are the log Bayes factor (lBf) from the comparison of the two models and the dashed lines indicate the classification thresholds at −1.1 in favor of the random model and 1.1 in favor of the GTR model.

**Figure S10:**
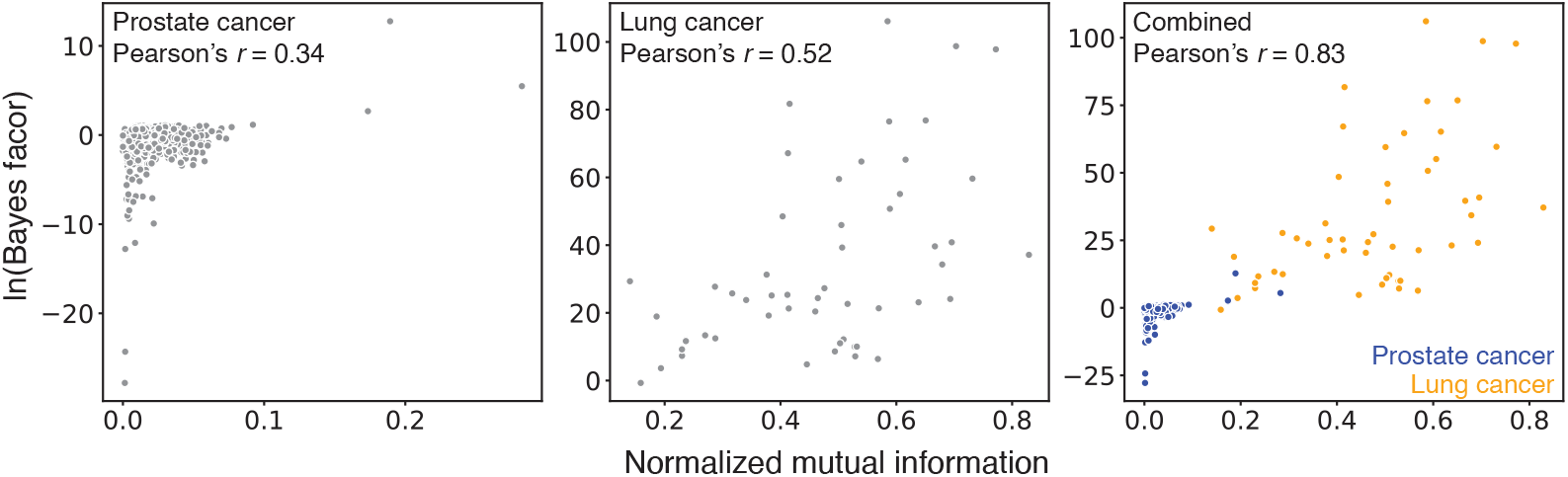
Relationship between the log Bayes factor from the random vs. GTR information content hypothesis test in **Fig. 4A** (*y*-axis) and the normalized mutual information (*x*-axis) computed from tissue transition count matrices across BEAM posterior distributions (see **Materials & Methods**). The prostate cancer data is shown on the left, the lung cancer data in the center, and the combined data on the right.

**Figure S11:**
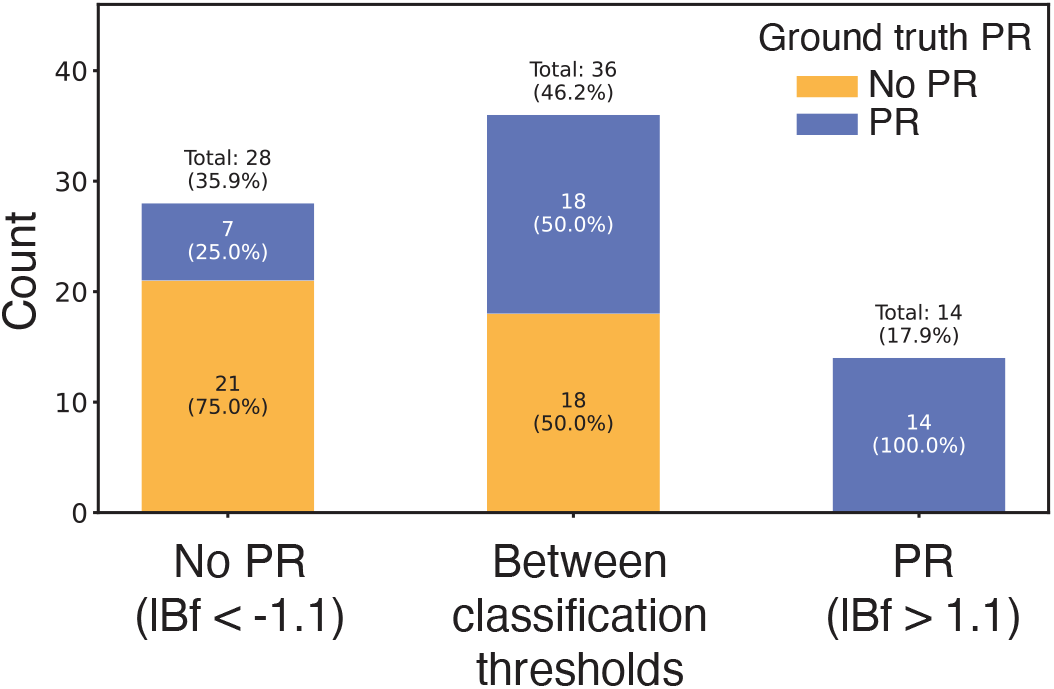
Simulated data were classified using a hypothesis test comparing primary reseeding (PR) and no PR models as done for real data in **Fig. 4B**. Simulations were selected from the larger variable-rates simulated dataset in **Fig. 2C** to include equal numbers of ground truth migration graphs with and without PR. True labels are indicated by color in the legend. Classification results are based on the log Bayes factor (lBF) where simulations are labeled positive for PR if lBF > 1.1, negative for PR if lBF < −1.1, or between classification thresholds if −1.1 < lBF < 1.1.

## Supplementary Tables

**Table S1:** The number of CPs with a transition from left lung (LL), right lung (RL), or mediastinum (M) to Liv at different edgewise posterior-probability thresholds. Counts are out of a total of 35 CPs with the Liv tissue observed.

| Migration Event | Consensus Threshold |  |  |  |  |  |  |
| --- | --- | --- | --- | --- | --- | --- | --- |
|  | 0.5 | 0.6 | 0.7 | 0.8 | 0.9 | 0.95 | 0.99 |
| LL → Liv | 13 | 10 | 8 | 4 | 2 | 0 | 0 |
| M → Liv | 27 | 27 | 25 | 25 | 19 | 16 | 9 |
| RL → Liv | 13 | 13 | 10 | 9 | 4 | 2 | 2 |

**Table S2:** The initial number of total CPs per mouse (left), the number of CPs per mouse selecting the GTR model in the random vs. GTR hypothesis test of data information in **Fig. 4A** (middle), and the number of CPs per mouse that selected the primary reseeding (PR) model in the no PR vs. PR hypothesis test in **Fig. 4B** (right) out of those that previously selected the GTR model (middle).

| **Mouse** | **Initial count** | **GTR selected** | **Primary reseeding selected** |
| --- | --- | --- | --- |
| MMUS1469 | 111 | 0 | 0 |
| MMUS1457 | 24 | 3 | 1 |
| MMUS1544 | 71 | 1 | 0 |
| MMUS1495 | 105 | 0 | 0 |
| MMUS1466 | 17 | 0 | 0 |
| MMUS1467 | 26 | 0 | 0 |
| MMUS1492 | 9 | 0 | 0 |
| MMUS1874 | 23 | 0 | 0 |
| MMUS1875 | 16 | 0 | 0 |
| MMUS1588 | 19 | 0 | 0 |
| **Overall count** | 421 | 4 | 1 |

